# Statistical evaluation of character support reveals the instability of higher-level dinosaur phylogeny

**DOI:** 10.1101/2023.01.25.525612

**Authors:** David Černý, Ashley L. Simonoff

## Abstract

The interrelationships of the three major dinosaur clades (Theropoda, Sauropodomorpha, and Ornithischia) have come under increased scrutiny following the recovery of conflicting phylogenies by a large new character matrix and its extensively modified revision. Here, we use tools derived from recent phylogenomic studies to investigate the strength and causes of this conflict. Using both the original and rescored dataset, we examine the global support for alternative hypotheses as well as the distribution of phylogenetic signal among individual characters. We find the three possible ways of resolving the relationships among the main dinosaur lineages (Saurischia, Ornithischiformes, and Ornithoscelida) to be statistically indistinguishable and supported by nearly equal numbers of characters in both matrices. While the changes made to the revised matrix increased the mean phylogenetic signal of individual characters, this amplified rather than reduced their conflict, resulting in greater sensitivity to character removal or coding changes and little overall improvement in the ability to discriminate between alternative topologies. We conclude that early dinosaur relationships are unlikely to be resolved without fundamental changes to both the quality of available datasets and the techniques used to analyze them.

---

The relationships between the three major clades that comprise Dinosauria (Theropoda, Sauropodomorpha, and Ornithischia) have historically been uncertain. In his seminal work that coined the name Ornithischia for herbivorous dinosaurs characterized by an opisthopubic pelvis, Seeley (1887) regarded this taxon as only distantly related to the theropods and sauropodomorphs, which he grouped together as Saurischia (Fig. 1). This view of dinosaur polyphyly later extended to Saurischia itself (Romer, 1956; Charig et al., 1965; Reig, 1970; Thulborn, 1975; Charig, 1982), leaving the ornithischians, theropods, and sauropodomorphs as three lineages independently descended from non-dinosaurian archosaurs of uncertain interrelationships.

**Fig. 1:**
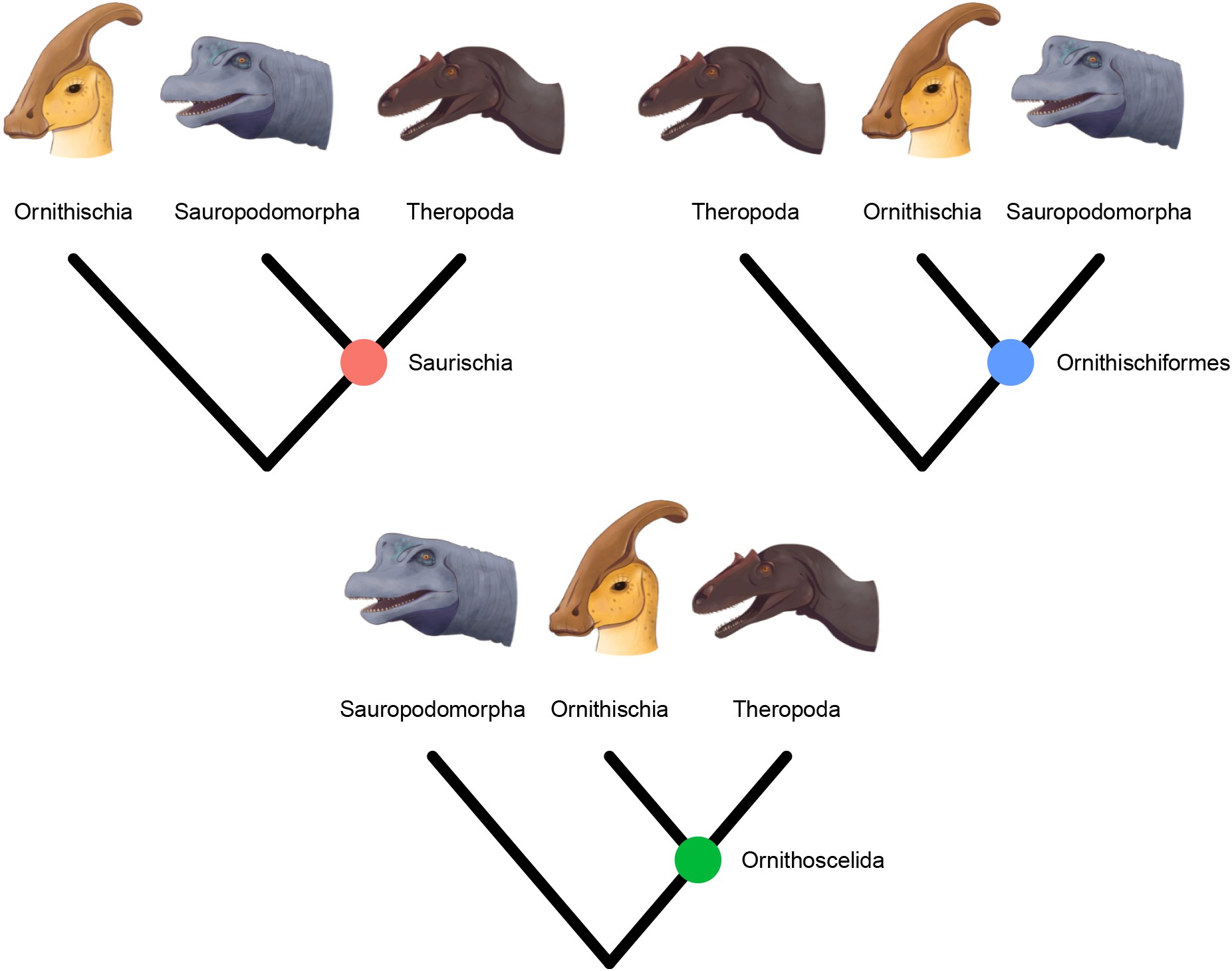
Three alternative hypotheses of early dinosaur phylogeny evaluated in this study.

The recognition of dinosaur monophyly failed to immediately clarify the relationships among the three clades, as the earliest arguments in favor of a monophyletic Dinosauria were coupled not only with the continued use of the Saurischia–Ornithischia dichotomy, but also with the seemingly contradictory suggestion that the ornithischians may have evolved from “prosauropods” (= early sauropodomorphs) (Bakker and Galton, 1974; Bonaparte, 1976; Cooper, 1981). In the mid-1980s, the latter scenario was formalized as a phylogenetic hypothesis linking Ornithischia and Sauropodomorpha to the exclusion of Theropoda (Paul, 1984, 1988; Sereno, 1984) in a group variously termed Ornithischiformes (Cooper, 1985) or Phytodinosauria (Bakker, 1986) (Fig. 1). However, this hypothesis was abandoned after the first rigorous application of algorithmic phylogenetics to nonavian dinosaurs by Gauthier (1986), who provided detailed character evidence uniting the theropods and sauropodomorphs into a monophyletic Saurischia, cementing a view of dinosaur phylogeny that would remain uncontested for the following three decades (Novas, 1996; Benton, 2004; Nesbitt, 2011).

Using a large new dataset, Baron et al. (2017a; henceforth BEA) recently destabilized this view by lending support to the third possible (and previously unforeseen) way of resolving the branch in question, namely, a clade formed by Theropoda and Ornithischia to the exclusion of Sauropodomorpha, for which the authors adopted Huxley’s (1870) name Ornithoscelida (Fig. 1). Seven months later, Langer et al. (2017; henceforth LEA) published a response using a rescored version of BEA’s character matrix with 9 taxa added, which recovered the traditional Saurischia–Ornithischia dichotomy, albeit with virtually no statistical support. The resulting controversy surrounding early dinosaur phylogeny has not been resolved by subsequent attempts to further rescore or add supposed key taxa such as *Pisanosaurus* (Baron et al., 2017b; Baron, 2019) and *Chilesaurus* (Baron and Barrett, 2017; Müller et al., 2018; Baron and Barrett, 2018; Müller and Dias-da Silva, 2019), nor by the use of more sophisticated phylogenetic methods such as time-free Bayesian inference (Parry et al., 2017) and Bayesian tip-dating (Griffin et al., 2022).

Despite the considerable interest generated by BEA’s contribution, there have been few attempts to quantify the relative support for each of the candidate topologies, distinguish between low information content and internal conflict, or identify the characters that may drive such conflict – lines of investigation that are more typical of phylogenomics (Reddy et al., 2017; Shen et al., 2017; Pease et al., 2018) than morphological phylogenetics. LEA took early steps in this direction by demonstrating that when their data was analyzed in a parsimony framework, the reinstated Saurischia hypothesis was statistically indistinguishable from either of the alternatives (Langer et al., 2017). However, subsequent studies have mostly reverted to reporting a single point estimate of the phylogeny, without testing whether its support significantly exceeded that of the next best hypothesis. The occasional attempts to use the number of extra steps relative to the most parsimonious tree for this purpose (Baron et al., 2017a; Baron and Barrett, 2017) suffer from the fact that this quantity has no statistical interpretation (Felsenstein, 2004). Moreover, despite the acknowledged centrality of character scoring differences to the conflict among the resulting topologies (Langer et al., 2017; Baron et al., 2017b; Müller et al., 2018; Baron and Barrett, 2018), only one study to date has attempted to determine which rescorings drove the difference between the hypotheses of early dinosaur phylogeny favored by BEA’s and LEA’s datasets (Goloboff and Sereno, 2021), and its methodological scope was limited to parsimony.

Here, we use statistical tools drawn from phylogenomics to evaluate the relative support for the three hypotheses of large-scale dinosaur phylogeny in the BEA and LEA datasets, and to conduct detailed assessments of character support. We demonstrate the presence of pervasive conflict both across each dataset as a whole and among the subsets of characters with the strongest phylogenetic signal. We further show that although LEA’s extensive changes to BEA’s character coding dramatically altered the distribution of phylogenetic signal across the matrix, they did little to help discriminate among the three alternative topologies, which remain indistinguishable both before and after LEA’s recoding. Our results suggest that there are many more plausible hypotheses of early dinosaur phylogeny than usually acknowledged, and that selecting between them may be beyond the reach of current character matrices and the techniques used to analyze them. We provide recommendations pertaining to both data-related and methodological aspects of the problem, and conclude that care should be taken to properly account for the uncertainty surrounding higher-level dinosaur phylogeny in downstream analyses.

## Results

### Maximum likelihood analyses

Our initial round of 100 maximum likelihood (ML) searches suggested that estimating early dinosaur phylogeny from the two datasets (Table 1) would present considerable difficulty due to the ruggedness of the resulting tree space. The high difficulty scores (BEA = 0.596, LEA = 0.649) were mostly driven by a lack of topological congruence, since each of the 100 ML estimates had a unique topology that slightly (average pairwise normalized Robinson-Foulds distance: BEA = 0.330, LEA = 0.395) but appreciably differed from those of the remaining trees (Supplementary Table 1). For the BEA dataset, the best-scoring tree was generally similar to the original parsimony estimate (Supplementary Fig. 1) and strongly supported a monophyletic Ornithoscelida (ultrafast bootstrap [UFBoot] = 99%). In contrast, the LEA dataset supported a version of the Ornithischiformes hypothesis which nested Ornithischia within Sauropodomorpha (Supplementary Fig. 2). The sister-group relationship between ornithischians and derived sauropodomorphs again received substantial support (UFBoot = 97%), while the broader clade uniting Ornithischiformes proper with early sauropodomorphs was less robust (UFBoot = 94%).

**Table 1:** Properties of the phylogenetic datasets analyzed in this study. The number of ordered characters is given as reported by BEA and LEA. For LEA, the proportion of missing data also includes the 109 polymorphic codings present in the original matrix, which are functionally equivalent to an unknown state under maximum likelihood.

|  | BEA | LEA |
| --- | --- | --- |
| Taxa | 74 | 83 |
| Characters | 457 | 457 |
| Ordered | 39 | 39 |
| Constant | 6 | 9 |
| Binary | 375 | 374 |
| Of those autapomorphic: | 45 | 43 |
| Of those parsimony-informative: | 330 | 331 |
| Multistate | 76 | 74 |
| Of those autapomorphic: | 2 | 1 |
| Of those parsimony-informative: | 74 | 73 |
| Proportion missing | 58.01% | 55.58% |

Trees that were constrained to alternative early dinosaur topologies (Saurischia and Ornithischiformes for BEA, Saurischia and Ornithoscelida for LEA) exhibited log-likelihoods that fell well within the range yielded by the initial 100 unconstrained ML searches, suggesting that the three hypotheses were statistically indistinguishable for both datasets. We formally corroborated this result using multiple likelihood-based topology tests, all of which indicated that none of the three topologies was significantly better or worse than the others (Tables 2, 3). The topological constraints were accommodated by the two datasets in markedly different ways. When their monophyly was enforced, Saurischia and Ornithischiformes were subtended by near-zero-length branches in the trees produced by the BEA dataset (Supplementary Figs. 3, 4), indicating a lack of characters that could be convincingly interpreted as saurischian or ornithischi form synapomorphies. For the LEA dataset, enforcing the monophyly of Saurischia yielded a topology that closely resembled the original parsimony estimate (cf. Supplementary Fig. 5 and Fig. 1 of Langer et al., 2017), while the tree optimized under an Ornithoscelida constraint showed an idiosyncratic topology that nested Ornithischia deep within theropods (Supplementary Fig. 6), reminiscent of hypotheses recently proposed by Baron (2019) to account for stratigraphic incongruence in ornithischian origins.

**Table 2:** Relative fit of the three alternative early dinosaur topologies to the BEA dataset. Best (unconstrained) and constrained ML trees are listed in order of decreasing likelihood (*L*). The bootstrap proportion (bp) from the resampling estimated log-likelihood (RELL) method is shown along with the *p*-values from the Kishino-Hasegawa (KH), Shimodaira-Hasegawa (SH), weighted Shimodaira-Hasegawa (wSH), and approximately unbiased (AU) tests, as well as the confidence value (*c*) based on expected likelihood weight (ELW). Significant rejection of (—) or the failure to reject (+) a given topology are indicated next to each test statistic.

| Hypothesis | ln *L* | bp (RELL) | *p* (KH) | *p* (SH) | *p* (wSH) | *p* (AU) | *c* (ELW) |
| --- | --- | --- | --- | --- | --- | --- | --- |
| Ornithoscelida | -6368.110126 | 0.754 + | 0.826 + | 1 + | 0.946 + | 0.824 + | 0.748 + |
| Saurischia | -6377.249639 | 0.162 + | 0.174 + | 0.26 + | 0.254 + | 0.218 + | 0.163 + |
| Ornithischiformes | -6377.484497 | 0.0844 + | 0.114 + | 0.247 + | 0.194 + | 0.222 + | 0.0888 + |

**Table 3:** Relatie fit of the three alternative early dinosaur topologies to the LEA dataset. Best (unconstrained) and constrained ML trees are listed in order of decreasing likelihood. Abbreviations as in Table 2.

| Hypothesis | $\ln L$ | bp (RELL) | $p$ (KH) | $p$ (SH) | $p$ (wSH) | $p$ (AU) | $c$ (ELW) |
| --- | --- | --- | --- | --- | --- | --- | --- |
| Ornithischiformes | -7023.11918 | 0.723 + | 0.882 + | 1 + | 0.962 + | 0.832 + | 0.718 + |
| Saurischia | -7034.61052 | 0.102 + | 0.118 + | 0.29 + | 0.195 + | 0.234 + | 0.106 + |
| Ornithoscelida | -7035.848825 | 0.175 + | 0.181 + | 0.273 + | 0.246 + | 0.208 + | 0.177 + |

### Character-wise support

In both datasets, the distribution of character support for each of the three hypotheses was indistinguishable from uniform, both across the matrix as a whole (multinomial test; BEA: *p* = 0.063, LEA: *p* = 0.668) and within most of the individual anatomical partitions (Fig. 2b,d). Despite the overall similarity of character support distributions between the two matrices, we found substantial differences at the level of individual characters. Only 70 out of the 447 overlapping non-constant characters ranked the three hypotheses in the same way in terms of their log-likelihoods, less than the one-sixth expected by chance. Similarly, only 145 characters preferred the same hypothesis, a number that is also indistinguishable from the one-third expected by chance (multinomial test: *p* = 0.726). While the number of characters supporting Ornithoscelida decreased as a result of LEA’s rescoring (from 174 to 147) and the number of characters supporting Saurischia marginally increased (from 141 to 143), even in the LEA dataset, more characters supported Ornithoscelida than Saurischia, although the difference was negligible (Fig. 2b,d).

**Fig. 2:**
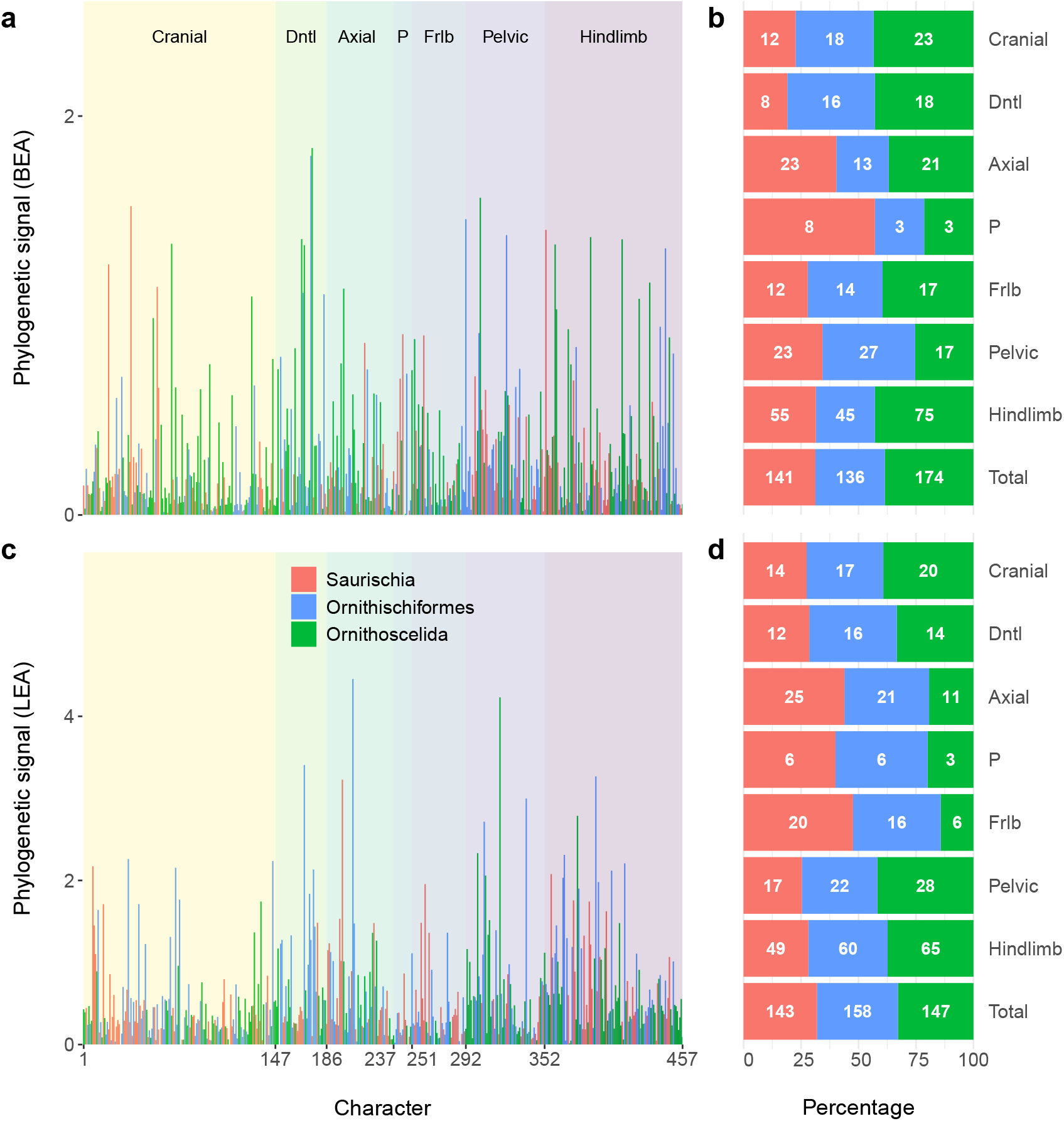
Distribution of phylogenetic signal across the two datasets. Phylogenetic signal of every nonconstant character from the BEA (a) and LEA (c) dataset is plotted against its placement in the matrix and colored by the preferred (highest-likelihood) hypothesis. The proportions and counts of characters preferring a given hypothesis are further shown for the BEA (b) and LEA (d) datasets as well as the individual anatomical regions by which they are organized. Abbreviations: Dntl = dental, P = pectoral, Frlb = forelimb.

Examining support for competing hypotheses in terms of simple preference (i.e., by recording which tree yields the highest log-likelihood score for a given character) fails to account for the fact that most characters do not strongly prefer any of the three alternatives, rendering the resulting log-likelihood differences negligible and overly sensitive to minor branch length differences. Indeed, the character-wise log-likelihood difference (ΔCLS) between the best and second best hypotheses exceeds a threshold of 0.5 for less than 10% of characters in the BEA dataset and 30% of characters in the LEA dataset, although we note that these proportions are still substantially higher than is typical of the phylogenomic datasets for which the threshold was originally defined (Shen et al., 2017; Francis and Canfield, 2020). In the BEA dataset, the majority of these “strong” characters (29 out of 45) support Ornithoscelida, which is also the case for a plurality (63 out of 137) of such characters in the LEA dataset.

To obtain a more fine-grained view of key characters driving the support for particular topologies, we employed the method suggested by Shen et al. (2017) and calculated the phylogenetic signal (PS) of each character as the mean of the absolute values of the three pairwise log-likelihood differences (Saurischia vs. Ornithischiformes, Saurischia vs. Ornithoscelida, Ornithischiformes vs. Ornithoscelida). Relative to the BEA dataset, the LEA dataset displays substantially higher mean (0.612 vs. 0.295) as well as maximum (4.454 vs. 1.840) phylogenetic signal values, consistent with its higher number of strong characters. The PS values of individual characters also differ considerably between the two datasets (Fig. 2a,c). In particular, among the characters with outlier PS values (more than three standard deviations above the mean), only one (character 169, serrations of maxillary and dentary teeth) is shared between the BEA matrix (in descending order of PS: 175, 174, 303, 37, 292, 353, 323, 387, 167, 411, 68, 360, 169, 444) and the LEA matrix (206, 318, 169, 391, 198, 338, 377, 306). Gradual removal of 1, 5, or 10 characters with the highest PS values or of all characters whose PS values represented outliers never caused either matrix to switch from the preferred topology to an alternative one (Supplementary Figs. 7–14). The exclusion of high-PS characters made little difference to the high statistical support for Ornithoscelida in the BEA dataset (UFBoot = 96–99%; Supplementary Figs. 7–10) but caused a drastic erosion of support for Ornithischiformes in the LEA dataset (Ornithischia + derived sauropodomorphs: UFBoot = 69–93%; Ornithischiformes proper + early sauropodomorphs: UFBoot = 52–79%; Supplementary Figs. 11–14).

To further distinguish among different ways in which a character can attain a high PS value, we separately compared each pair of hypotheses (Fig. 3), focusing on those characters that also emerged as outliers in at least two of the three pairwise comparisons. These corresponded to characters that strongly favored a particular hypothesis, strongly disfavored one, or both. The results reveal substantial conflict among the high-PS characters, as each matrix contains characters that both strongly favor and strongly disfavor its globally preferred topology (Ornithoscelida for BEA, Ornithischiformes for LEA) as well as one or both of its alternatives (Fig. 3, Table 4). Neither of the characters that strongly favor Ornithoscelida in the BEA dataset was among the 21 synapomorphies of this clade originally identified by Baron et al. (2017a), although one such synapomorphy (character 360, state 1: medial bowing of the femur forming a gentle curve) was found to strongly disfavor Saurischia in the present analysis (Table 4). Similarly, there is virtually no overlap between the characters strongly favoring or disfavoring one of the three hypotheses in the present study, and the “keystone” characters recently identified using a parsimony-based approach (Goloboff and Sereno, 2021).

**Fig. 3:**
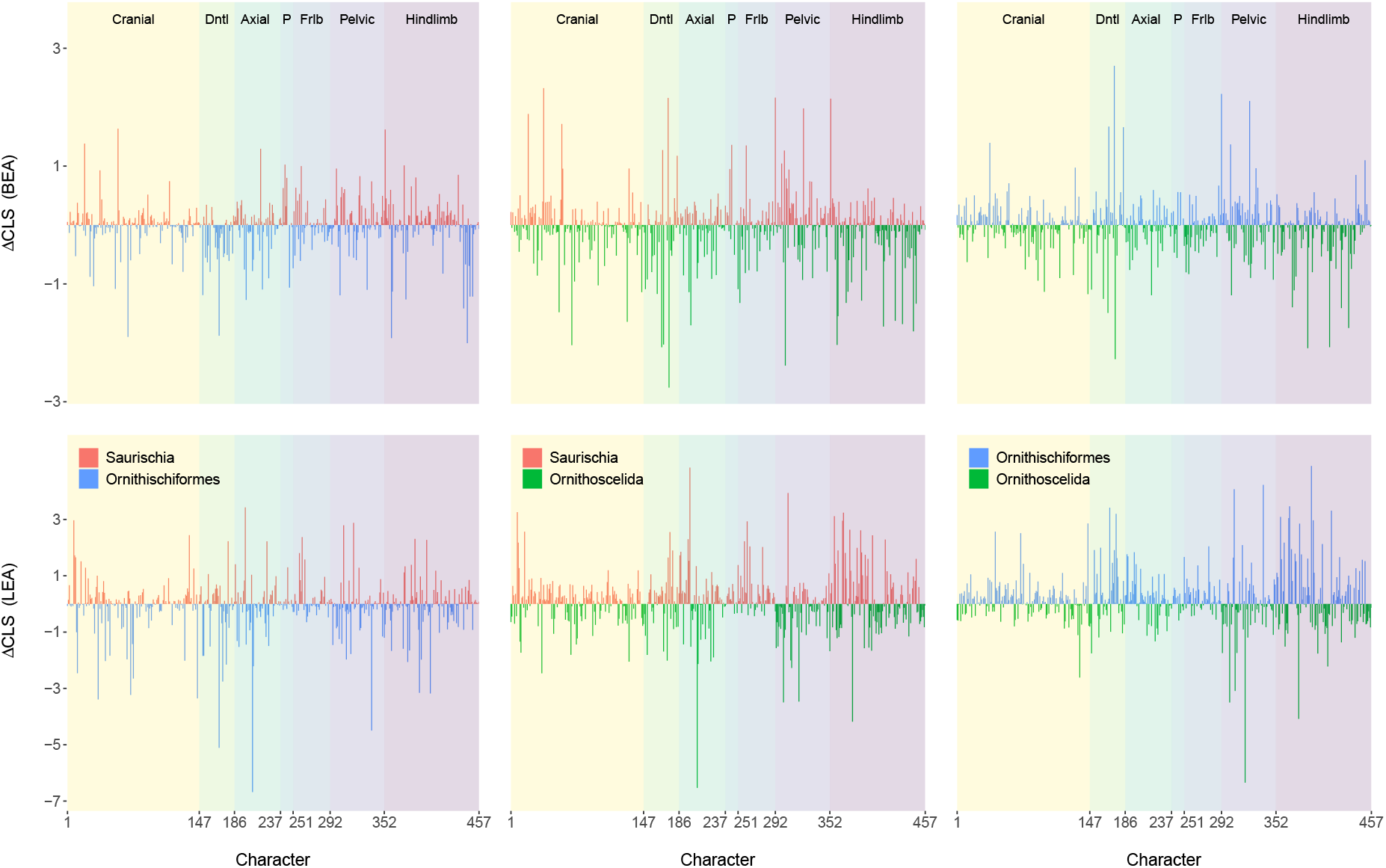
Pairwise comparisons of character support for the three alternative early dinosaur topologies. For each pair of hypotheses, the difference in character-wise log-likelihood values (△CLS) of every nonconstant character from the BEA (top) and LEA (bottom) datasets is plotted against its placement in the matrix. Anatomical region abbreviations as in Fig. 2.

**Table 4:** Outlier characters in the BEA and LEA matrices. Of = Ornithischiformes, Os = Ornithoscelida, S = Saurischia. Characters were only included if both their phylogenetic signal (PS) and at least two of the three pairwise log-likelihood differences [△CLS(S, Of), △CLS(S, Os), △CLS(Of, Os)] were more than three standard deviations above the mean. Hypotheses that are strongly favored (+) or disfavored (–) relative to both alternatives are indicated next to each character.

| Character | PS in BEA | PS in LEA | Outlier in BEA | Outlier in LEA |
| --- | --- | --- | --- | --- |
| 68 | 1.360 | 0.352 | S (–) | — |
| 167 | 1.383 | 0.409 | Os (+) | — |
| 169 | 1.352 | 3.407 | S (–) | Of (+) |
| 174 | 1.801 | 1.275 | Os (–) | — |
| 175 | 1.840 | 0.103 | Os (+) | — |
| 198 | 0.139 | 3.228 | — | S (+) |
| 206 | 0.605 | 4.454 | — | S (–) |
| 292 | 1.483 | 0.193 | Os (–) | — |
| 306 | 0.262 | 2.716 | — | Os (–) |
| 318 | 0.370 | 4.231 | — | Os (+), Of (–) |
| 323 | 1.403 | 0.500 | Os (–) | — |
| 338 | 0.493 | 2.998 | — | Of (+) |
| 353 | 1.429 | 0.738 | S (+) | — |
| 360 | 1.356 | 1.112 | S (–) | — |
| 377 | 0.057 | 2.787 | — | Os (+) |
| 391 | 0.226 | 3.267 | — | Of (+) |
| 444 | 1.337 | 0.780 | S (–) | — |

### Rescoring of individual characters

To determine which of LEA’s coding changes most contributed to the difference between the topologies yielded by the original and rescored datasets, we successively replaced the scoring of each character in either matrix by its scoring from the opposite matrix, and checked whether this change was sufficient for the re-estimated ML tree to switch to a different topology (Fig. 4). The procedure had little impact on the BEA dataset’s support for Ornithoscelida, which proved robust to the rescoring of any one of the 438 applicable characters and remained high on average (mean UFBoot = 98.8%; Fig. 4a). Only three characters (110, 114, 387) caused the support for Ornithoscelida to drop below the 95% threshold when changed to their scorings in the LEA dataset. Of these, none was optimized as an ornithoscelidan synapomorphy in BEA’s original analysis (Baron et al., 2017a), and while all favored Ornithoscelida over the alternatives in the original BEA matrix, only character 387 did so strongly (ΔCLS > 0.5 relative to the next best hypothesis). In the LEA matrix, all three characters ranked Saurischia and Ornithoscelida as the best- and worst-supported hypothesis, respectively, but the log-likelihood difference between the two exceeded the 0.5 threshold only for character 387.

**Fig. 4:**
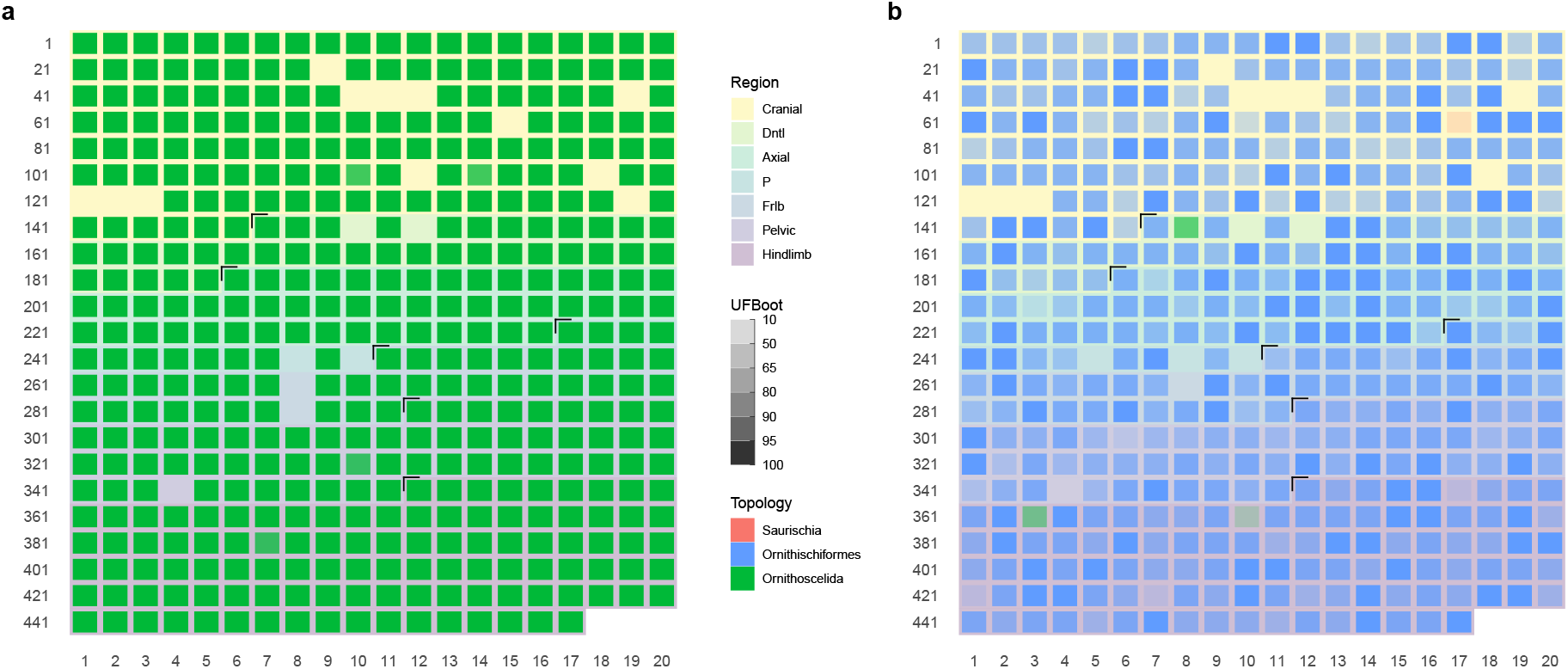
Support for alternative topologies after rescoring each matrix one character at a time. Characters changed from BEA’s to LEA’s coding (a) or from LEA’s to BEA’s coding (b) are arranged by row following their placement in either dataset; black marks denote the boundaries between adjacent anatomical regions denoted by different background colors. Color and opacity of individual heatmap cells denote the topology of the ML tree inferred after rescoring the character in question and the ultrafast bootstrap (UFBoot) support for that topology’s focal clade, respectively. Gaps correspond to characters whose coding did not differ between the two matrices or characters that would be rendered constant by having their coding changed to that of the opposite matrix. Anatomical region abbreviations as in Fig. 2.

In contrast, reverting four characters to their original scoring in the BEA matrix proved sufficient to flip the ML result from the LEA dataset (Ornithischiformes) either to Ornithoscelida (characters 148, 363, 370) or to Saurischia (character 77) (Fig. 4b), producing highly idiosyncratic topologies (Supplementary Figs. 15–18). In the ornithoscelidan trees, Ornithischia was deeply nested within theropods (with *Panguraptor* and *Zupaysaurus* consistently recovered closer to Ornithischia than to other theropods), similar to the results obtained from a constrained analysis of the unmodified LEA matrix (Supplementary Fig. 6). The support for the nodes uniting Ornithischia with its successive theropod outgroups was generally low, although one such node received a UFBoot value of 97% in the analysis based on rescoring character 370 (Supplementary Fig. 18), consistent with the fact that state 2 of this character (prominent, wing-like anterior trochanter) represented an ornithoscelidan synapomorphy under BEA’s original scoring (Baron et al., 2017a; see also Baron and Barrett, 2017). The only saurischian topology found was similarly unconventional and poorly supported (Supplementary Fig. 15). None of the characters that caused a switch from Ornithischiformes to Ornithoscelida favored Ornithischiformes in the LEA dataset, and only two of them (characters 148 and 370) favored Ornithoscelida in the BEA dataset. Notably, character 77 consistently ranked Saurischia as the worst of the three hypotheses under both BEA and LEA scorings; the fact that its rescoring was nevertheless sufficient for Saurischia to emerge as the preferred hypothesis indicates an extreme degree of instability across the LEA matrix. This is also borne out by the fact that even among those trees which continued to support Ornithischiformes, altering the scoring of a single character caused the UFBoot support for this clade to drop from 97% to (on average) 89.5% (Fig. 4b). Only 105 of the 441 applicable characters (23.8%) upheld Ornithischiformes with UFBoot support greater than 95% upon rescoring.

## Discussion

By repurposing a protocol originally developed for phylogenomic data (Shen et al., 2017), we found that both the BEA and LEA datasets are unable to conclusively resolve the interrelationships of major dinosaur clades. All three hypotheses of overall dinosaur phylogeny – Saurischia, Ornithischiformes, and Ornithoscelida – remain plausible, and neither dataset shows any of these to be significantly better or worse than the alternatives (Tables 2, 3). Our results suggest that this is not due to low information content of the two matrices; in fact, the proportion of characters that strongly discriminate between the best and second best hypothesis (ΔCLS > 0.5) is far higher (10–30%) than typical for phylogenomic data (0.1–7%; Shen et al., 2017; Francis and Canfield, 2020). Similarly, although both matrices exhibit a highly uneven distribution of phylogenetic signal and contain several outlier character strongly favoring or disfavoring particular topologies, the results were not exclusively driven by a handful of such outliers, since their removal had limited impact on the support for one topology or another (Supplementary Figs. 7–14). Instead, we hypothesize that the lack of meaningful statistical support for any of the three hypotheses (partially obscured by high UFBoot values) is due to pervasive conflict among individual characters. Limiting the focus to the characters with the highest PS values revealed patterns of conflict (Table 4) that were similar to those observed across each matrix as a whole (Fig. 2), explaining why their removal did little to change the underlying distribution of support.

According to almost every metric employed in this study, the LEA dataset produces less stable results than the BEA dataset. It yields a higher phylogenetic difficulty score (Supplementary Table 1), exhibits a more uniform distribution of character support for the three hypotheses (Fig. 2c, d), and is more sensitive to the exclusion of outlier characters (Supplementary Figs. 11–14) despite containing fewer of them (Table 4). The LEA matrix was also less robust to changes in character coding, as demonstrated by its tendency to flip the ML tree from one topology to another after reverting a single character to its original scoring by BEA (Fig. 4). Consistent with the findings above, this greater degree of instability was not caused by weaker phylogenetic signal in the LEA matrix. In fact, although LEA’s rescorings and taxon additions only marginally improved the completeness of the matrix (Table 1), they resulted in a markedly higher mean character-wise phylogenetic signal (Supplementary Fig. 19), and amplified the log-likelihood differences between competing hypotheses both for the dataset as a whole (Table 3) and for individual characters (Supplementary Fig. 20). In effect, the stronger phylogenetic signal present in the recoded and expanded matrix only served to amplify, rather than eliminate, underlying conflict within the dataset. In light of the failure to resolve this rampant conflict by extensive coding changes, we outline several alternative recommendations for identifying its sources and assessing the relative support for competing topologies in its presence.

First, we recommend re-examining the original dataset at a deeper level, as the recovery of divergent yet statistically indistinguishable topologies may serve to highlight fundamental issues in the underlying character data. Poorly formulated characters should ideally be redefined rather than simply rescored. For example, LEA’s coding changes that caused character 174 (recurvature of maxillary and dentary teeth) to lose the outlier status it originally had in the BEA dataset (Table 4) still took place in the framework of the vague definition inherited from BEA, who in turn modified it from an even earlier study (Butler et al., 2008). The character description provides no quantitative criterion for differentiating between teeth possessing strong, weak, or no recurvature, allowing for more or less arbitrary coding changes. Indeed, the scoring of *Efraasia* was changed by LEA from no recurvature to weak recurvature without explicit justification or photographic evidence, and on the basis of the same published sources which BEA cited in support of their own original coding. The problem of vague character descriptions and subjective scoring decisions is widespread in the two matrices (e.g., characters 114, 216, 266, 337) and compounded by a number of additional issues, some of which were noted by BEA and LEA themselves. These include multiple instances of scoring taxa for characters that cannot be ascertained from their known material (Baron et al., 2017b) and character non-independence (Langer et al., 2017). While the former issue represents a fundamental problem with the data and a potentially serious biasing factor, especially when it stems from coding taxa based on assumed rather than observed morphologies (Giribet, 2010; Gee, 2021), the latter might be mitigated at the methodological level. Frameworks capable of dealing with hierarchical and correlated characters are under development (Tarasov, 2022), as are ontology-based methods for their semi-automated detection and characterization (Eliason et al., 2019; Porto et al., 2022). More advanced methods may ultimately also alleviate other problems with existing matrices. The BEA and LEA datasets contain instances of problematic state delimitation that make it impossible to assign certain morphologies to any existing state (e.g., character 28, length of antorbital fenestra equal to 10–15% of skull length), or merge distinct morphologies into a single state (e.g., character 77, state 0: paraquadratic foramen small or absent). Both exemplify a more general problem, namely the arbitrary discretization of characters that would be more naturally treated as continuous (Poe and Wiens, 2000). This ubiquitous feature of morphological phylogenetic datasets reflects long-standing methodological limitations, which have only recently been overcome by phylogenetic software packages that simultaneously implement models for discrete and continuous characters, making it possible to combine both types of data in a single analysis (Höhna et al., 2016; Bouckaert et al., 2019).

Second, instead of attempting to rescore or redefine an entire dataset in an indiscriminate “shotgun” approach, it is prudent to determine which characters are responsible for the signal in that dataset’s results. While LEA re-examined the putative ornithoscelidan synapomorphies identified by BEA at an admirable level of detail (Langer et al., 2017), both previous findings (Goloboff and Sereno, 2021) and our own results demonstrate that the characters supporting a particular topology do not always coincide with the characters that map as the synapomorphies of that topology’s focal clade. Methods for identifying such critical characters are now available in parsimony (Goloboff and Sereno, 2021), maximum-likelihood (Shen et al., 2017; Francis and Canfield, 2020), and Bayesian (Porto et al., 2022) frameworks, and can be profitably used to narrow the focus of potential rescoring efforts, which often involve the time- and labor-intensive process of gathering data from first-hand observations of multiple museum specimens. The benefits of comprehensively revising a pre-existing character matrix have to be weighed against the costs inherent to such an effort, as well as the risk of introducing new errors into the data. In morphological phylogenetics, extensive reuse and iterative expansion of pre-existing matrices gives rise to complicated dataset genealogies (Gee, 2021; Regalado Fernández and Werneburg, 2022) in which coding error can propagate and compound over time. On the other hand, our results suggest that the effort invested into the comprehensive rescoring of a large pre-existing dataset can be difficult to justify: the three hypotheses of early dinosaur phylogeny were statistically indistinguishable based on the original BEA dataset, and remain such after LEA’s extensive revision of it.

Third, we urge paleontologists to quantify the uncertainty associated with their phylogenetic hypotheses using well-characterized tools with a clear statistical interpretation. When alternative ways of resolving a given branch are of interest, as in the controversy surrounding early dinosaur phylogeny, we encourage the community to move beyond the mere reporting of a phylogenetic point estimate toward explicitly testing it against the next best alternative. Following recent practice (Wu et al., 2023), we used a variety of likelihood ratio tests (LRTs) to this end (Shimodaira and Hasegawa, 1999; Shimodaira, 2002; Strimmer and Rambaut, 2002), but other approaches are possible. Bayesian inference differs from maximum likelihood in its treatment of nuisance parameters such as branch lengths or the parameters of the substitution and rate heterogeneity models, which are jointly optimized with the parameter of interest (topology) in maximum likelihood but marginalized over in Bayesian methods (Huelsenbeck et al., 2002). Both approaches have their advantages (Holder and Lewis, 2003), and as a result, Bayes factors – a Bayesian equivalent of LRTs, relying on marginal rather than joint likelihoods (Kass and Raftery, 1995) – can represent a useful alternative way of evaluating competing topologies (Suchard et al., 2005; Bergsten et al., 2013). Although most of the best-performing marginal likelihood estimators are much more computationally demanding than joint likelihood inference (Fourment et al., 2019), the minute size of phylogenetic datasets employed by paleontologists makes their application relatively easy, and Bayes factor topological comparisons are accordingly starting to see use in dinosaur phylogenetics (O’Connor et al., 2020).

Taken together, our results suggest that large-scale dinosaur phylogeny is much more poorly understood than commonly acknowledged. In particular, despite the widespread perception that saurischian monophyly is challenged primarily by the recently proposed Ornithoscelida hypothesis (Felice et al., 2020; Müller and Garcia, 2020; Castiglione et al., 2022), the earlier Ornithischiformes hypothesis receives comparable support, and is in fact weakly preferred when the LEA dataset is analyzed using maximum likelihood (Table 3). Additional support for Ornithischiformes was also recently detected in an independent dataset (Baron, 2022). Moreover, not only the three major hypotheses – Saurischia, Ornithischiformes, and Ornithoscelida – but also a number of their variations nesting the ornithischians deep within Sauropodomorpha or Theropoda (Supplementary Figs. 2, 6, 16–18) cannot be ruled out at present. Indeed, the specific variation on the Ornithischiformes topology recovered in this study shows the ornithischians to be not just sister to, but rather nested within Sauropodomorpha, with a clade of Carnian to early Norian sauropodomorphs (approximately corresponding to the Guaibasauridae of Ezcurra, 2010 or the Saturnaliidae of Langer et al., 2019) branching off before the ornithischians. This result is consistent with a phylogeny inferred from the LEA dataset by Parry et al. (2017) using time-free Bayesian inference, showing that the two model-based methods yield topologies that are much more similar to each other than to those favored by parsimony. By positing an early divergence of the early Late Triassic sauropodomorphs, this scenario helps reduce the temporal gap between the first appearance of Ornithischia and its sister group (Baron, 2019), and shows remarkable congruence with the early suggestions that the ornithischians may have arisen from within “prosauropods” (Bakker and Galton, 1974; Bonaparte, 1976; Cooper, 1981; Bakker, 1986).

Our findings suggest that higher-level dinosaur interrelationships represent a phylogenetic problem of considerable difficulty that is unlikely to be conclusively resolved by minor additions and superficial modifications to the datasets currently in use. While our investigation was limited to two such datasets (Baron et al., 2017a; Langer et al., 2017), there are few reasons to believe that other matrices currently employed by dinosaur paleontologists are free of the problems identified here. Indeed, the main alternative to the datasets examined here has repeatedly lent support to yet another nonstandard hypothesis nesting the putatively non-dinosaurian Silesauridae within Ornithischia (Cabreira et al., 2016; Müller and Garcia, 2020; Norman et al., 2022), indicating that the number of plausible early dinosaur phylogenies proliferates even further when not only the three major clades, but also species-poor lineages such as Herrerasauridae and Silesauridae are taken into consideration. As a result, the increasingly common practice of repeating comparative analyses under the Saurischia and Ornithoscelida topologies (Felice et al., 2020; Castiglione et al., 2022; Hendrickx et al., 2022) most likely severely understates the uncertainty associated with early dinosaur phylogeny, potentially leading to biased or overconfident conclusions.

## Methods

### Data

Our analyses were performed on the original character matrices used by BEA and LEA, whose properties are summarized in Table 1. Both were obtained from Graeme T. Lloyd’s database of previously pub-lished character matrices with standardized formatting (http://graemetlloyd.com/matrdino.html; last accessed May 1, 2022). Both BEA and LEA used *Euparkeria capensis* and *Postosuchus kirkpatricki* as outgroup taxa. The two datasets differed in 3366 scorings (10.0% of the total number of overlapping cells) when polymorphic codings were treated as distinct from missing data, and in 3350 scorings (9.9% of the total number of overlapping cells) when treating the two as equivalent (as in the maximum likelihood analyses). Only 4 of the 74 overlapping taxa (*Dromomeron gigas*, *Dromomeron gregorii*, *Dromomeron romerii, Postosuchus kirkpatricki*) and 10 of the 457 characters (50–52, 118, 121–123, 152, 250, 344) were unaffected by the changes. The much higher number of differences previously reported in the literature (8050 scorings, or 21.2% of the total number of LEA’s cells; Goloboff and Sereno, 2021) also reflects non-overlapping cells (representing the extra taxa added by LEA) as well as the notational distinction between missing data (“?”) and inapplicables (“-”), which has no analytical significance in currently used phylogenetic algorithms (but see Goloboff et al., 2021; Hopkins and St. John, 2021; Tarasov, 2022). Both datasets are organized by anatomical region, with cranial characters (1–146) followed by dental (147–185), axial (186–236), and pectoral (237–250) characters, and finally by characters pertaining to the forelimb (251–291), pelvis (292–351), and the hindlimb (352–457).

### Maximum likelihood analyses

We used maximum likelihood (ML) in our analyses of BEA’s and LEA’s datasets for several reasons. First, we found it useful to explore how the resulting topologies might change when using a parametric rather than nonparametric approach to phylogenetic inference, since method-dependent results may indicate the presence of within-dataset conflict (Betancur-R. et al., 2019). Second, despite its continued widespread use in the paleontological literature, maximum parsimony is well-known for its undesirable statistical properties compared to model-based methods (Felsenstein, 1978), including in the context of morphological phylogenetics (Wright and Hillis, 2014; O’Reilly et al., 2016, 2018; Puttick et al., 2019). Third, topologically constrained ML analyses allowed us to directly compare support between the three hypotheses using a number of well-established and easily interpretable frequentist tests.

All maximum likelihood analyses were performed using IQ-TREE v2.1.3 (Minh et al., 2020) with the default mix of starting trees (1 BioNJ + 99 parsimony trees). Both datasets were partitioned first into ordered and unordered characters and further by the number of character states, for a total of 6 partitions. In contrast to the number reported by the original authors (Table 1), only 36 and 37 characters from the BEA and LEA datasets were treated as ordered, respectively, since for characters 24, 334, and (for BEA) 180, only states 0 and 1 were observed. Branch lengths were treated as proportional among partitions (the -p command-line option in IQ-TREE), and unordered characters were assigned the M*k* model (Lewis, 2001) with *k* ranging from 2 to 5. Constant characters (BEA: 29, 59, 150, 245, 248, 268; LEA: 29, 59, 75, 112, 139, 150, 248, 268, 288) were excluded, and an ascertainment bias correction (Lewis, 2001) was applied to all partitions. Unlinked discrete gamma models of among-character rate heterogeneity were added to every substitution model except those applied to the character-poor unordered 5-state and ordered 4-state partitions.

### Difficulty assessment and tree searches

A number of methods have been developed to quantify the expected difficulty of phylogenetic analysis or the amount of data needed to resolve a particular node (Braun and Kimball, 2001; Poe and Chubb, 2004; Verbruggen et al., 2010); however, these often rely on models that are inapplicable to morphological evolution, such as the multispecies coalescent (Sayyari and Mirarab, 2018). To evaluate how challenging it would be to estimate early dinosaur interrelationships from the BEA and LEA datasets in a maximum-likelihood framework, we performed 100 topologically unconstrained tree searches on each dataset, and used the resulting ML trees to calculate a nonparametric difficulty measure recently proposed by Haag et al. (2022):

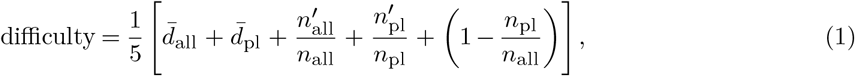

where 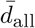 is the average pairwise normalized Robinson-Foulds (RF; Robinson and Foulds, 1981) distance between the *n*_all_ = 100 inferred trees, 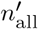 is the number of unique topologies among the 100 ML trees, 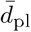 is the average pairwise normalized RF distance within a subset of plausible trees, *n*_pl_ is the number of trees included in this set, and 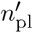 is the number of unique topologies present in the plausible set. Each term of Eq. (1) ranges from 0 to 1, as does the overall difficulty score equal to their unweighted mean. Depending on the resulting value, ML phylogenetic inference can range from trivial (0) to effectively impossible (1). Following Morel et al. (2020), the plausible tree set was constructed from all trees that were not found to be significantly worse than the best-scoring tree by any of the likelihood-based tests implemented in IQ-TREE. These were conducted using 10,000 approximate-bootstrap replicates (`-zb 10000 -zw -au`) generated by the resampling estimated log-likelihood method (Kishino et al., 1990) and included the Kishino-Hasegawa test (Kishino and Hasegawa, 1989), the unweighted and weighted SH test (Shimodaira and Hasegawa, 1999), the approximately unbiased test (Shimodaira, 2002), and expected likelihood weights (Strimmer and Rambaut, 2002). Postprocessing was carried out in the R statistical computing environment (R Core Team, 2021) using the packages *phytools* (Revell, 2012), *TreeTools* (Smith, 2019), and their respective dependencies.

After the difficulty assessment, we performed one more round of unconstrained ML searches consisting of 10 runs (each with 100 starting trees). The overall ML estimate was obtained by selecting the best-scoring tree from the pooled sample of the 100 exploratory and 10 final runs. To assess clade support, we additionally performed ultrafast bootstrap approximation (UFBoot) with 1000 replicates (-B 1000), either after the fact (if the best-scoring tree was found during the exploratory runs) or simultaneously with the main tree search (for the last 10 runs). In addition to being relatively robust to model misspecification, the ultrafast bootstrap is less biased than the standard nonparametric bootstrap (Hoang et al., 2017), and we also found it to be more numerically stable. Finally, we carried out topologically constrained analyses enforcing those hypotheses that were not supported by the unconstrained tree. Similar to the main analyses, each consisted of 10 runs performed simultaneously with 1000 UFBoot replicates.

### a. Character-wise support

We followed the protocol of Shen et al. (2017) to assess how support for the three competing hypotheses (S = Saurischia, Of = Ornithischiformes, Os = Ornithoscelida) was distributed across characters in both matrices, and to identify potential outliers. Using IQ-TREE, we estimated character-wise log-likelihood values for the ML trees yielded by both unconstrained and topologically constrained searches (`-wslr`). We first calculated the number of characters in either matrix that preferred a given hypothesis (i.e., yielded the least negative log-likelihood value under it), and repeated this calculation for individual anatomical partitions. Multinomial tests were carried out using the R package *EMT* (Menzel, 2021) to determine whether the resulting distributions were significantly different from uniform. We further determined the number of characters that ranked the three hypotheses in the same way in terms of their log likelihoods between the two matrices. Next, we calculated the phylogenetic signal (PS) of the *i*-th character (*C_i_*) in a given matrix following Eq. (5) of Shen et al. (2017):

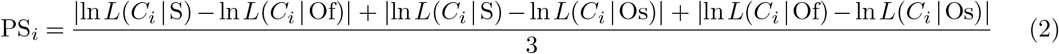

Since the observed distributions of PS values were heavier-tailed than those shown in Shen et al. (2017), we used a different criterion to identify outliers, defining them as those characters whose PS was more than three standard deviations above the mean (Francis and Canfield, 2020). To assess the influence exerted by characters with strong phylogenetic signal on the resulting early dinosaur topologies, we generated 8 subsampled datasets by removing 1, 5, and 10 characters with the highest PS values from either matrix, as well as those characters whose PS values represented outliers according to the above criterion (BEA: 14 characters, LEA: 8 characters). These subsampled matrices were then subject to unconstrained ML searches under the same settings as the original datasets (10 runs of 100 starting trees each + 1000 UFBoot replicates).

The definition of phylogenetic signal outlined above has recently been criticized for conflating cases in which one topology is strongly favored relative to either of the alternatives, and cases in which one topology is strongly disfavored relative to both alternatives that may nevertheless remain nearly indistinguishable from each other (Francis and Canfield, 2020). To tease apart these two scenarios, we further evaluated the difference in log-likelihood scores (ΔCLS) separately for each pair of hypotheses. For example, following Eq. (2) of Shen et al. (2017), the support lent to Saurischia over Ornithoscelida by the *i*-th character was calculated as:

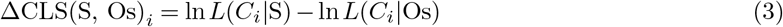

We then applied the same criterion for outliers to absolute ΔCLS values derived from each pairwise comparison, and identified those characters that represented outliers in at least two of the three comparisons. These corresponded to characters that either strongly favored or strongly disfavored a given topology relative to both of the alternatives. We also applied the criterion suggested by Francis and Canfield (2020) and identified those characters for which the log-likelihood difference between the best and second best hypotheses exceeded 0.5.

### Rescoring of individual characters

To identify characters whose rescoring may have had disproportionate impact on the resulting topology, we ran further ML analyses on modified versions of both datasets in which we successively recoded one character at a time to its scoring in the opposite dataset. We excluded from consideration those characters whose coding did not change between the two matrices (see “Data”) as well as those that would be rendered constant by reverting their scoring to that of the opposite dataset (see “Maximum likelihood analyses”), resulting in a total of 879 analyses. All of these were conducted under the same settings as the analyses of the original datasets (10 runs of 100 starting trees each + 1000 UFBoot replicates). Using a custom R script employing the package *phangorn* (Schliep, 2010), we scored each resulting ML topology for the recovery of Saurischia, Ornithischiformes, or Ornithoscelida, and extracted the UFBoot value of whichever of these three clades was present in the tree. To facilitate this process, we treated the names Saurischia, Ornithischiformes, and Ornithoscelida as referring to node-based clades, operationally defining them as (*Dilophosaurus* + *Plateosaurus*), (*Scelidosaurus* + *Plateosaurus*), and (*Dilophosaurus* + *Scelidosaurus*), respectively.

## Acknowledgments

We thank Graham J. Slater for valuable comments and advice, and to Catherine Wray for the illustrations used in Fig. 1. D.Č. further thanks Joseph F. Walker for the suggestion to apply phylogenomic methods of investigating site-wise support for alternative topologies to morphological datasets.

