## Supplementary Information for "Statistical evaluation of character support reveals the instability of higher-level dinosaur phylogeny"

| **Metric** | **BEA** | **LEA** |
|---|---|---|
| # of trees ($n_{\text{all}}$) | 100 | 100 |
| # of plausible trees ($n_{\text{pl}}$) | 67 | 51 |
| Avg. pairwise normalized RF distance, all trees ($\bar{d}_{\text{all}}$) | 0.3296 | 0.3952 |
| Avg. pairwise normalize RF distance, plausible trees ($\bar{d}_{\text{pl}}$) | 0.3203 | 0.3618 |
| # of unique topologies, all trees ($n'_{\text{all}}$) | 100 | 100 |
| # of unique topologies, plausible trees ($n'_{\text{pl}}$) | 67 | 51 |

**Supplementary Table 1:** Metrics computed to evaluate the difficulty of estimating higher-level dinosaur phylogeny from the Baron et al. (BEA) and Langer et al. (LEA) datasets. RF distance = Robinson-Foulds distance.

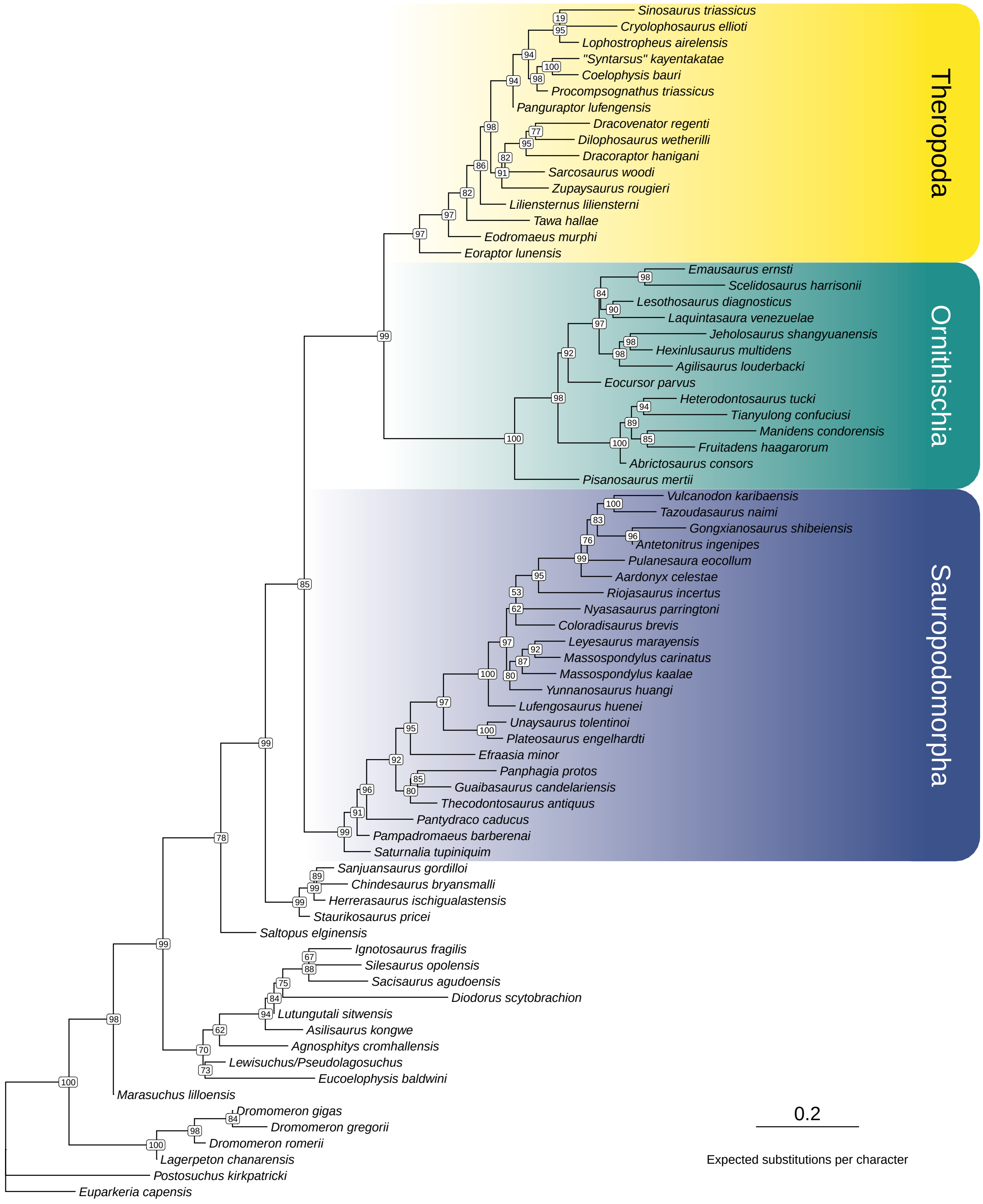


**Supplementary Fig. 1:** Unconstrained maximum likelihood tree inferred for the original BEA matrix ($\ln L = -6367.975$). Node labels denote ultrafast bootstrap values computed from 1000 replicates.

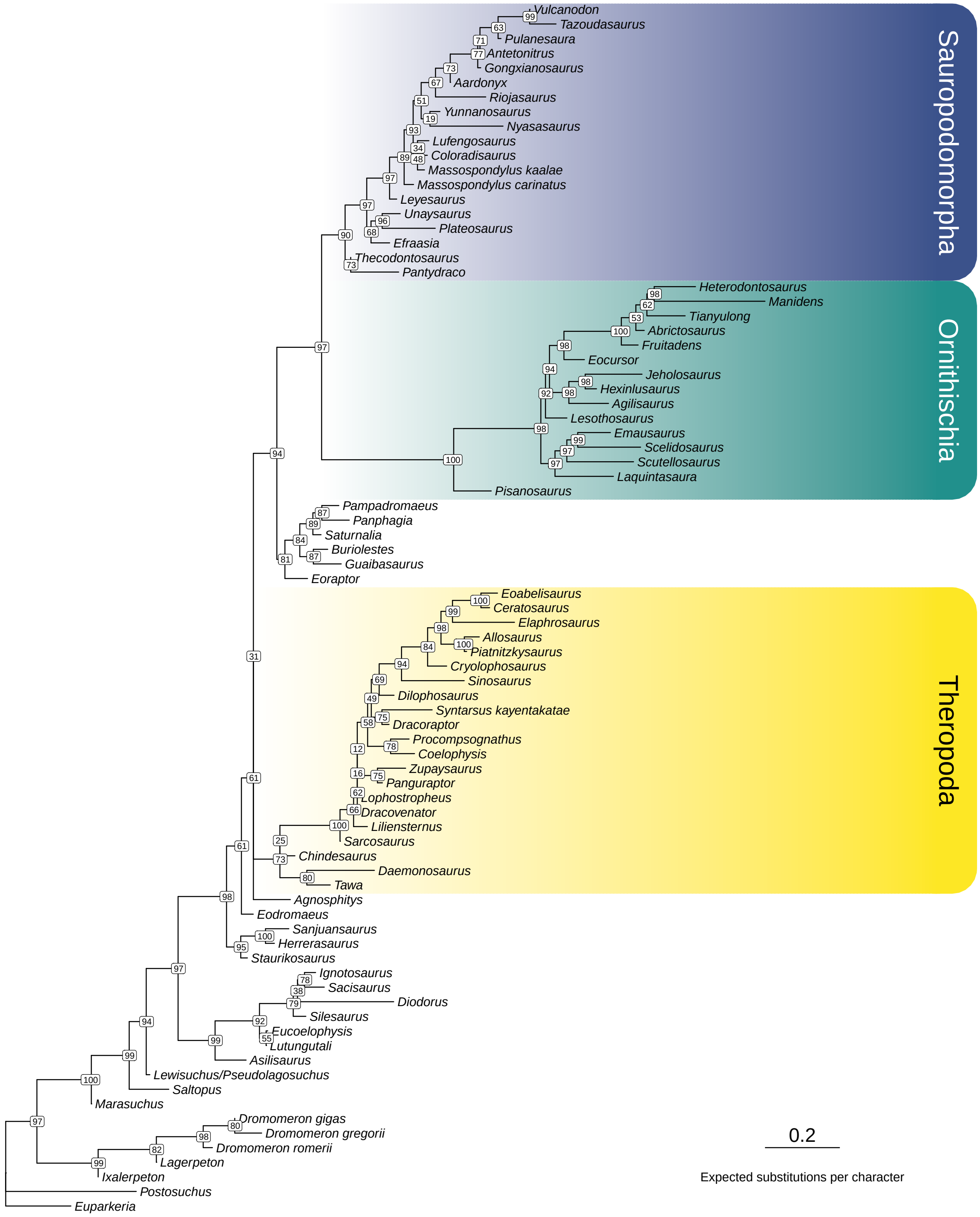


**Supplementary Fig. 2:** Unconstrained maximum likelihood tree inferred for the original LEA matrix ($\ln L = -7007.117$). Node labels denote ultrafast bootstrap values computed from 1000 replicates.

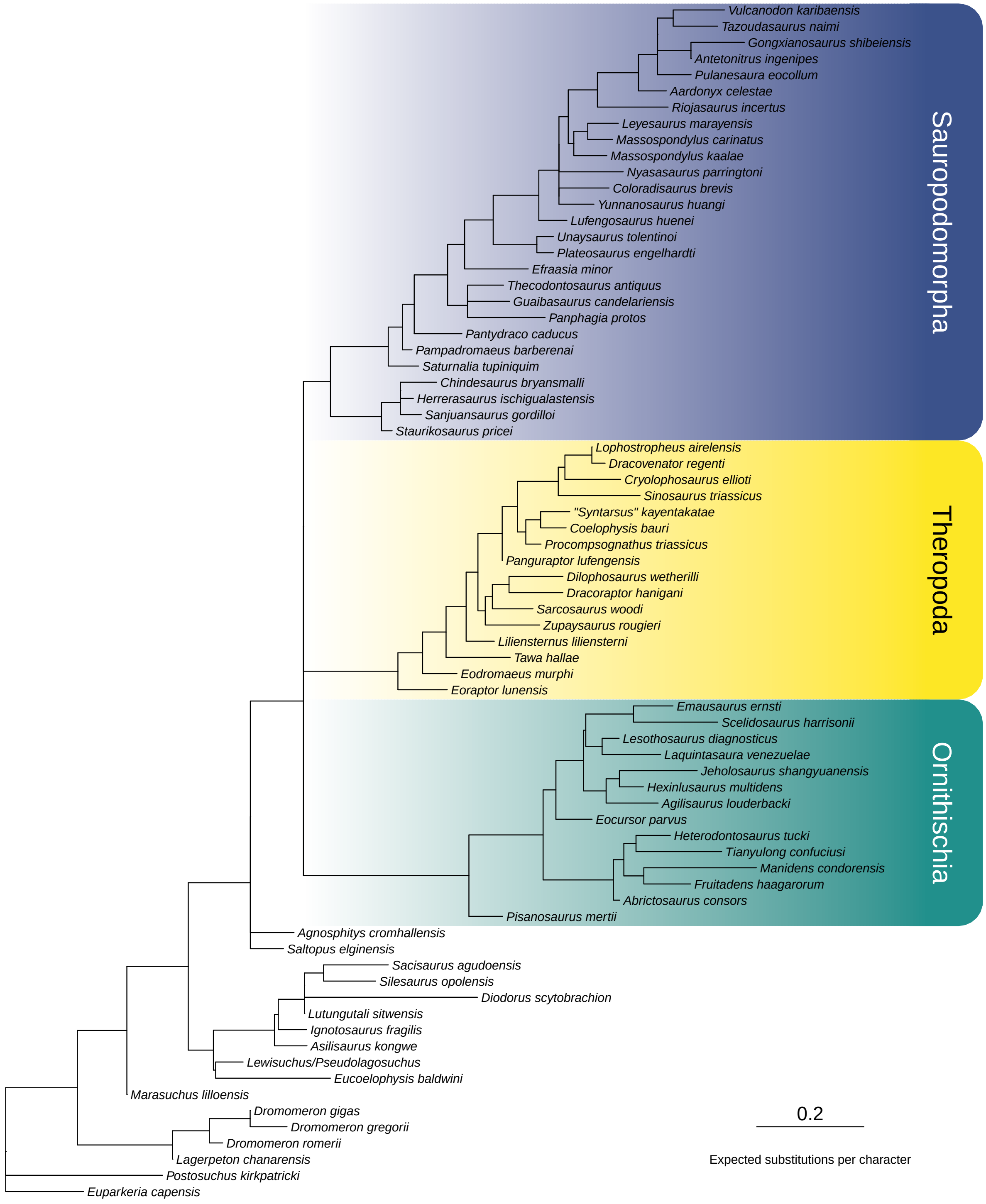


**Supplementary Fig. 3:** Maximum likelihood tree inferred for the original BEA matrix and constrained to recover Saurischia ($\ln L = -6376.854$).

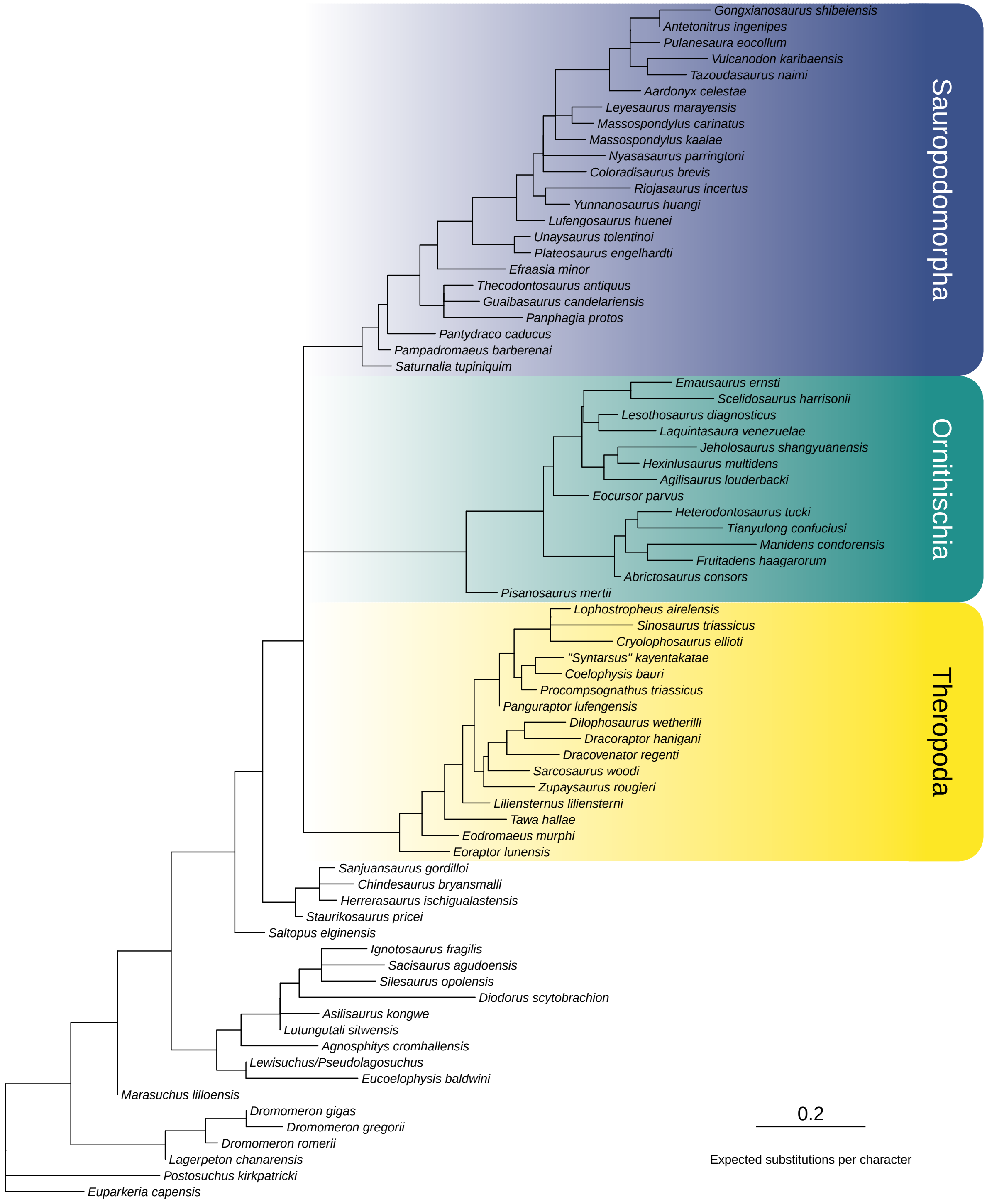


**Supplementary Fig. 4:** Maximum likelihood tree inferred for the original BEA matrix and constrained to recover Ornithischiformes (ln $L = -6377.425$).

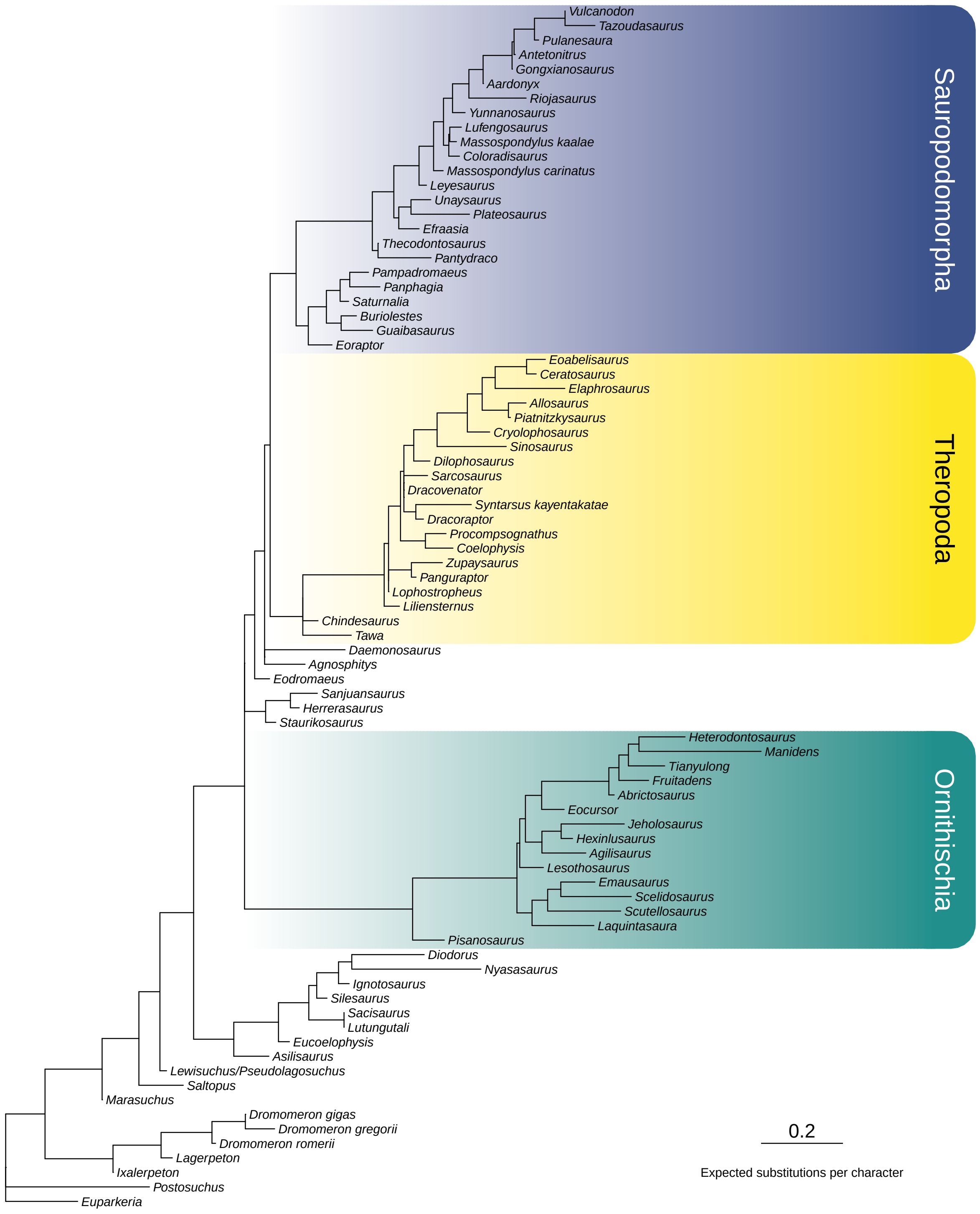


**Supplementary Fig. 5:** Maximum likelihood tree inferred for the original LEA matrix and constrained to recover Saurischia ($\ln L = -7017.246$).

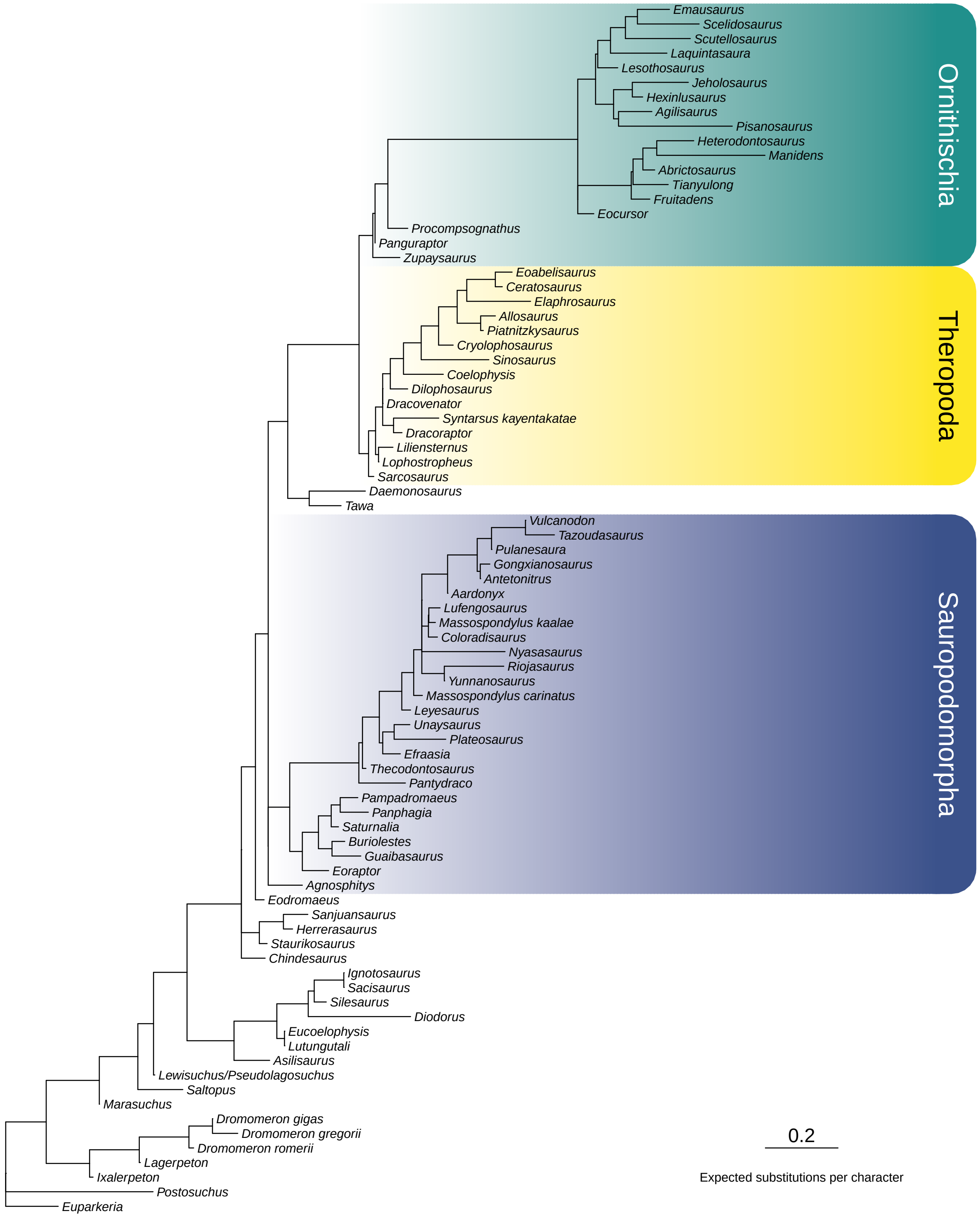


**Supplementary Fig. 6:** Maximum likelihood tree inferred for the original LEA matrix and constrained to recover Ornithoscelida ($\ln L = -7017.424$).

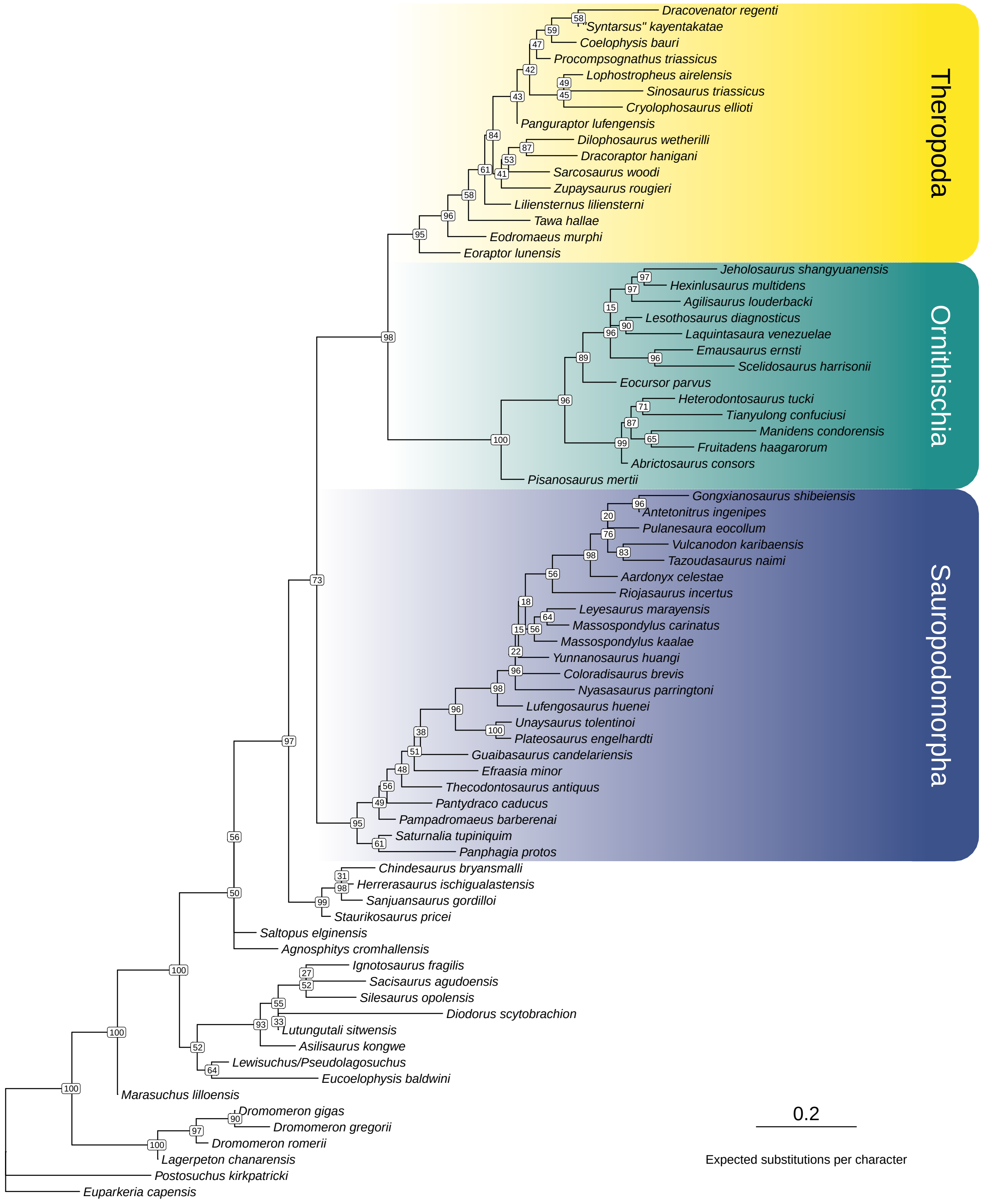


**Supplementary Fig. 7:** Maximum likelihood tree inferred for the BEA matrix without the single most decisive character (character 175; PS = 1.840) (ln $L = -6353.365$). Node labels denote ultrafast bootstrap values computed from 1000 replicates.

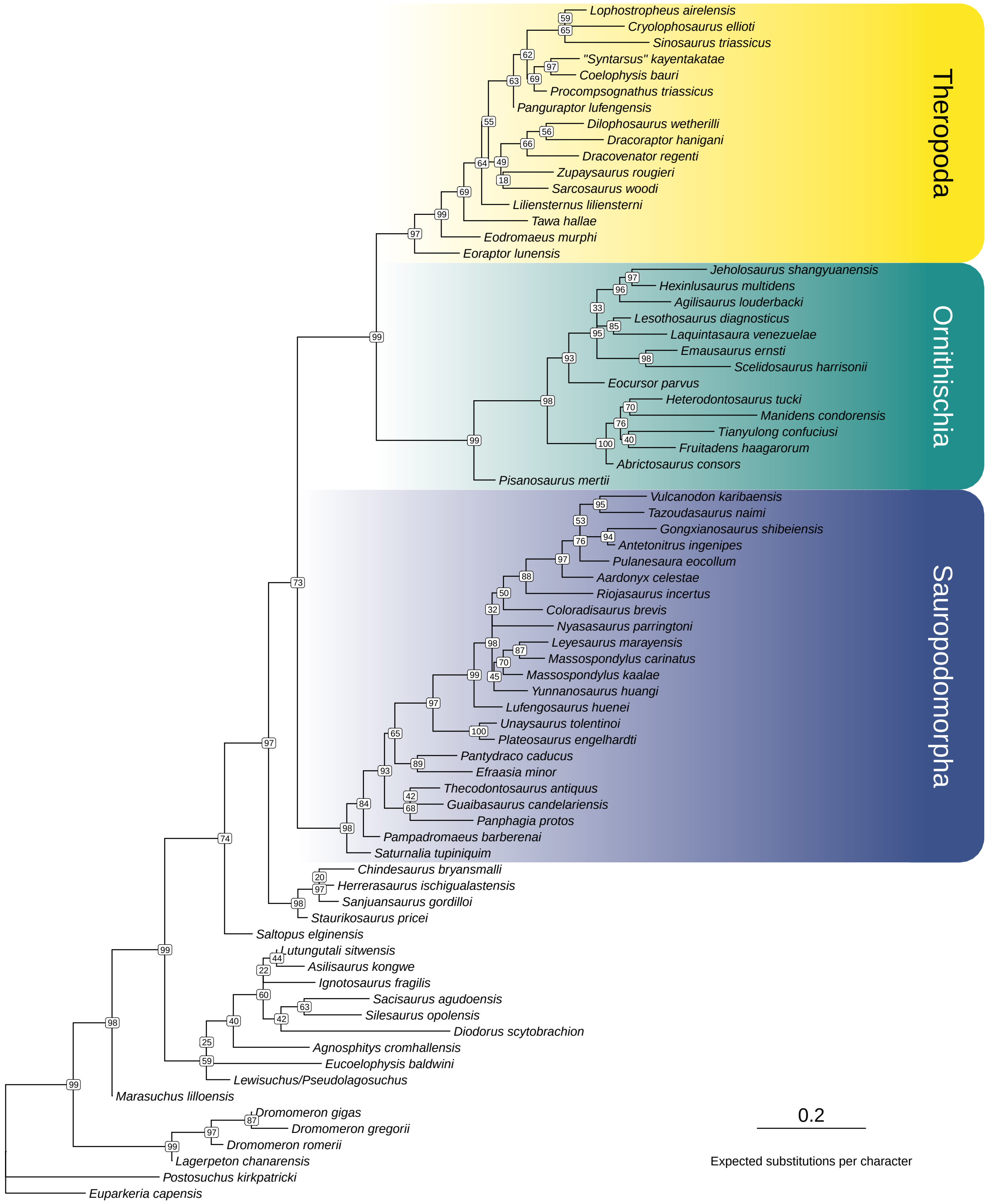


**Supplementary Fig. 8:** Maximum likelihood tree inferred for the BEA matrix without the 5 most decisive characters (characters 175, 174, 303, 37, 292; PS = 1.483–1.840) (ln $L = -6232.176$). Node labels denote ultrafast bootstrap values computed from 1000 replicates.

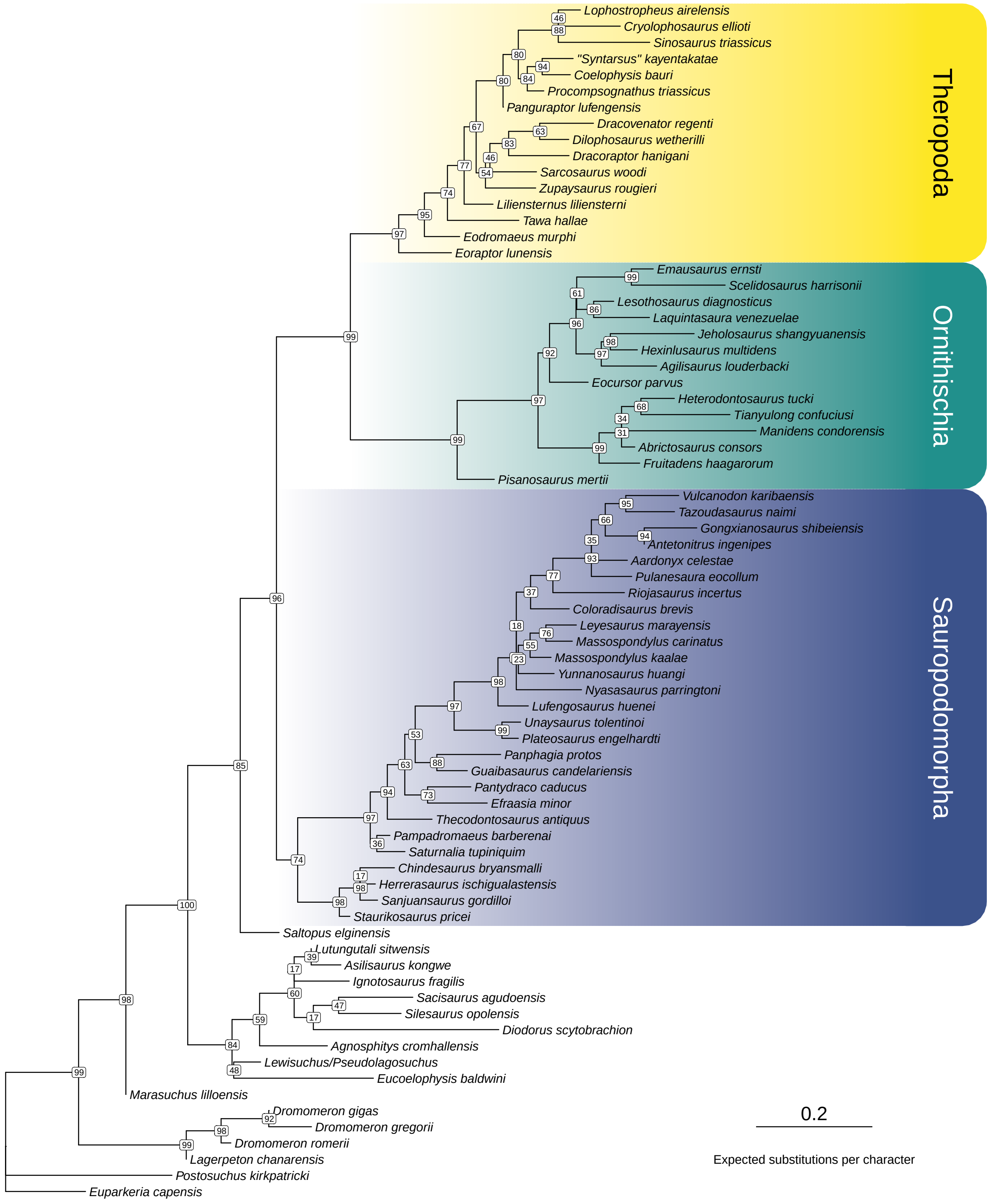


**Supplementary Fig. 9:** Maximum likelihood tree inferred for the BEA matrix without the 10 most decisive characters (characters 175, 174, 303, 37, 292, 353, 323, 387, 167, 411; PS = 1.382–1.840) ($\ln L = -6103.143$). Node labels denote ultrafast bootstrap values computed from 1000 replicates.

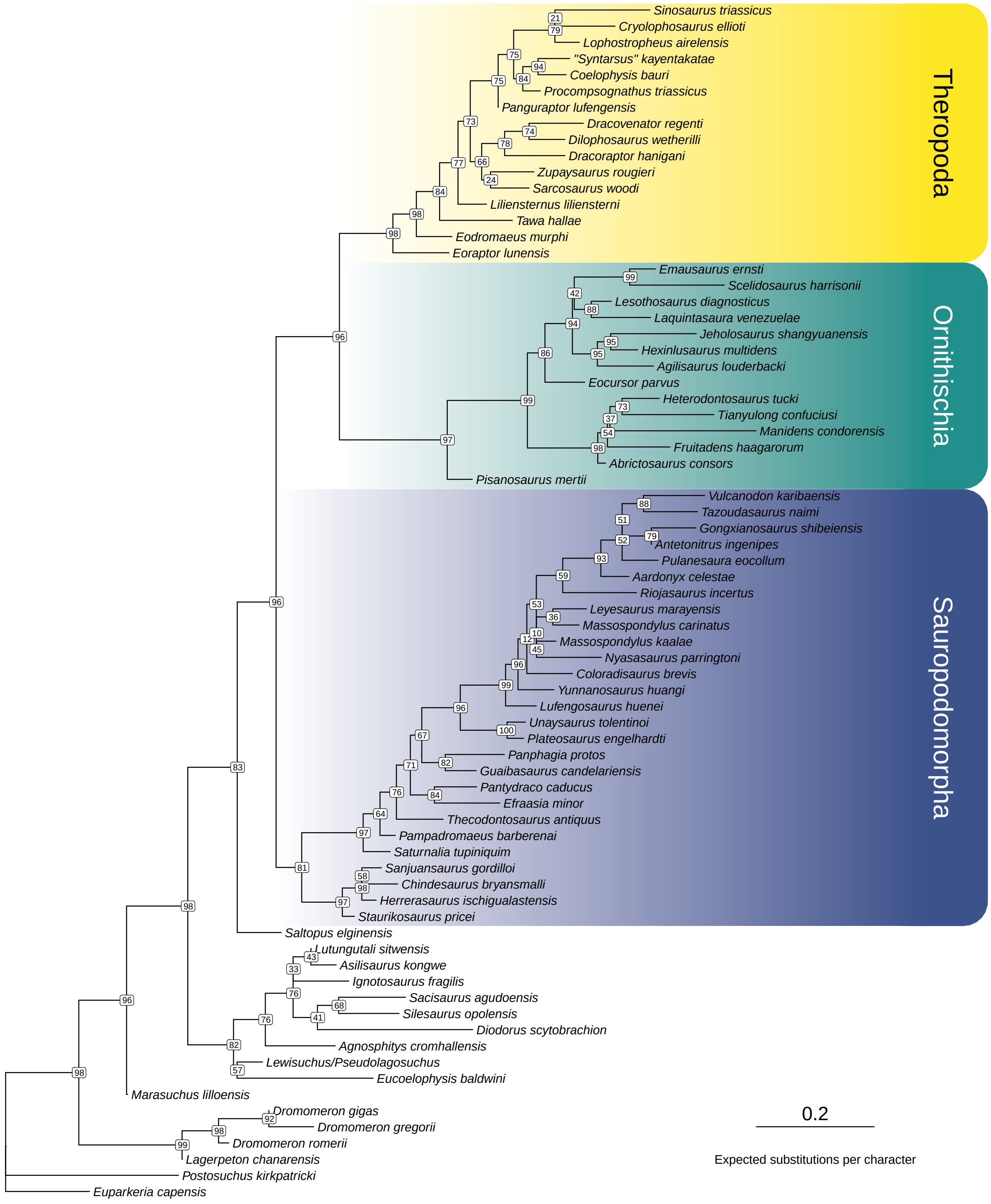


**Supplementary Fig. 10:** Maximum likelihood tree inferred for the BEA matrix without all characters with outlier PS values (characters 175, 174, 303, 37, 292, 353, 323, 387, 167, 411, 68, 360, 169, 444; PS = 1.337–1.840) (ln $L = -5992.320$). Node labels denote ultrafast bootstrap values computed from 1000 replicates.

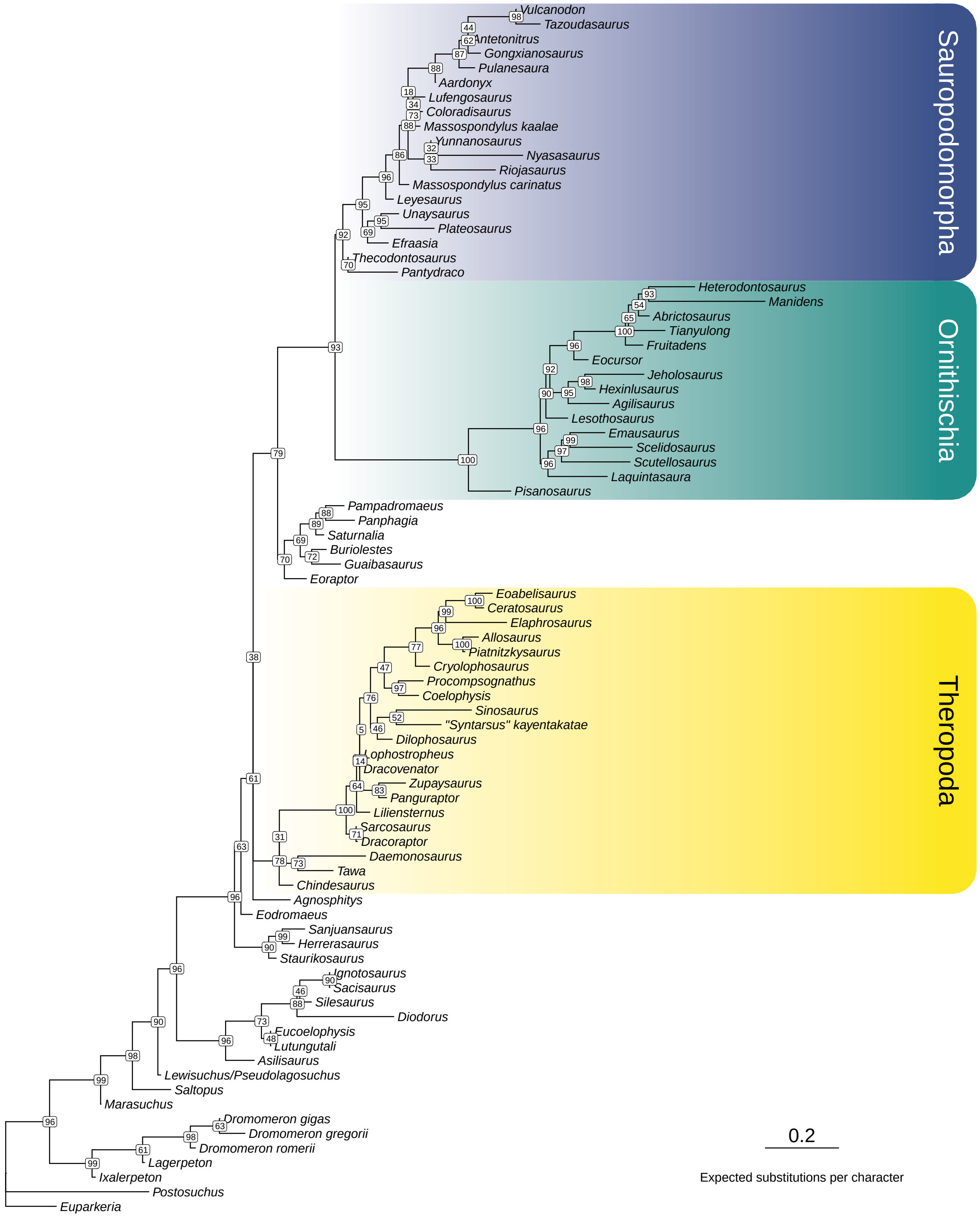


**Supplementary Fig. 11:** Maximum likelihood tree inferred for the LEA matrix without the single most decisive character (character 206; PS = 4.454) ($\ln L = -6972.146$). Node labels denote ultrafast bootstrap values computed from 1000 replicates.

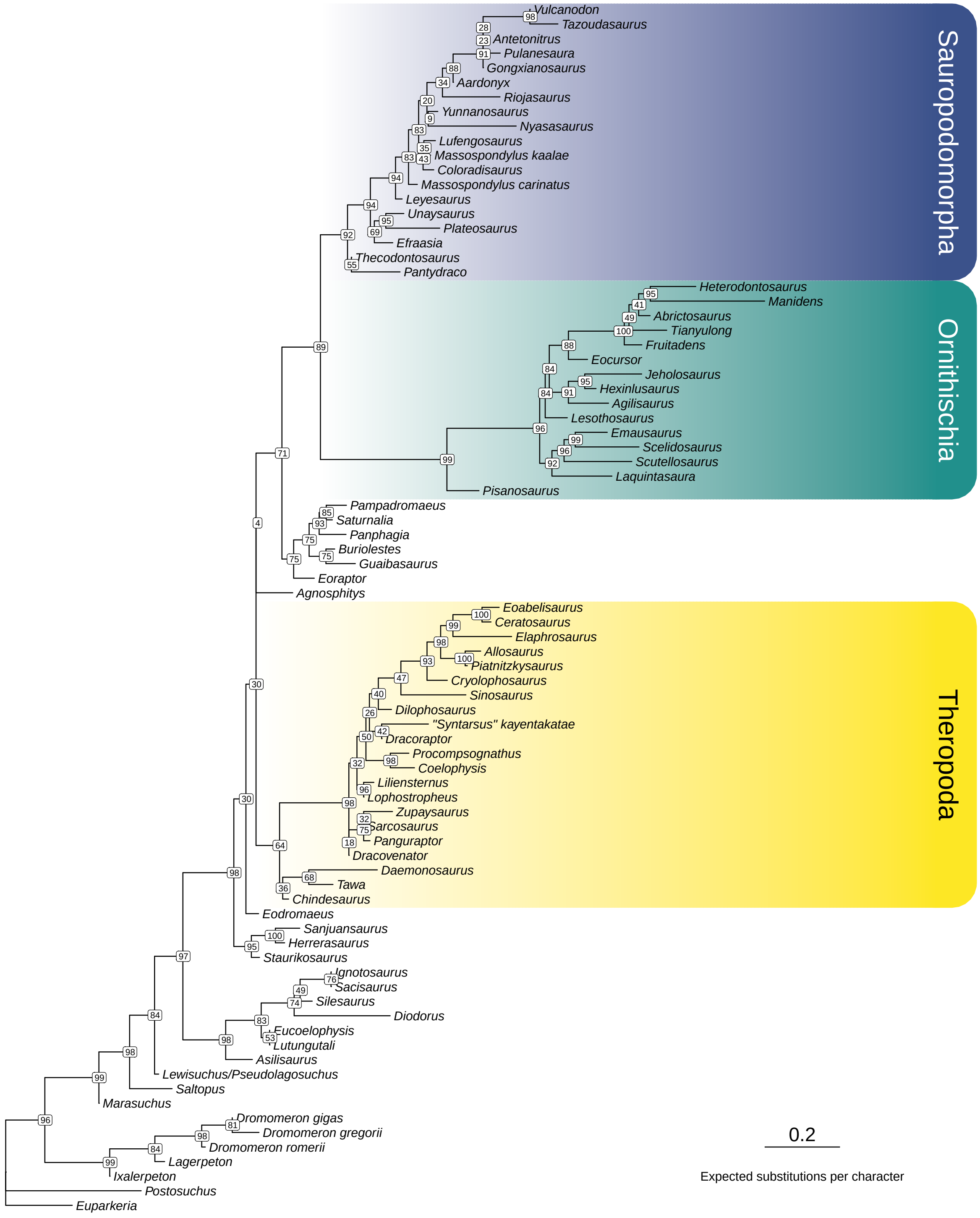


**Supplementary Fig. 12:** Maximum likelihood tree inferred for the LEA matrix without the 5 most decisive characters (characters 206, 318, 169, 391, 198; PS = 3.228–4.454) ($\ln L = -6869.597$). Node labels denote ultrafast bootstrap values computed from 1000 replicates.

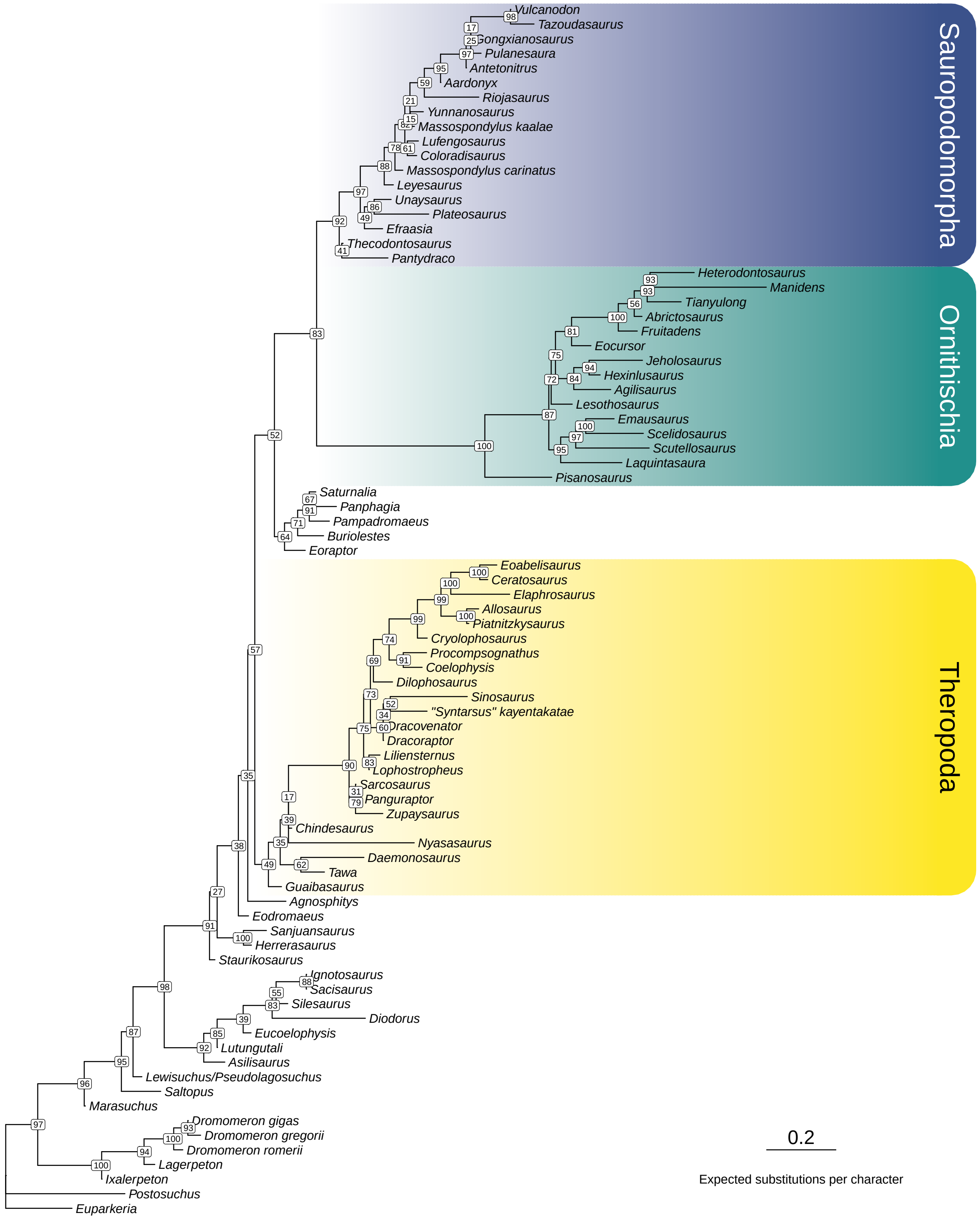


**Supplementary Fig. 13:** Maximum likelihood tree inferred for the LEA matrix without the 10 most decisive characters (characters 206, 318, 169, 391, 198, 338, 377, 306, 301, 367; PS = 2.311–4.454) (ln $L = -6795.273$). Node labels denote ultrafast bootstrap values computed from 1000 replicates.

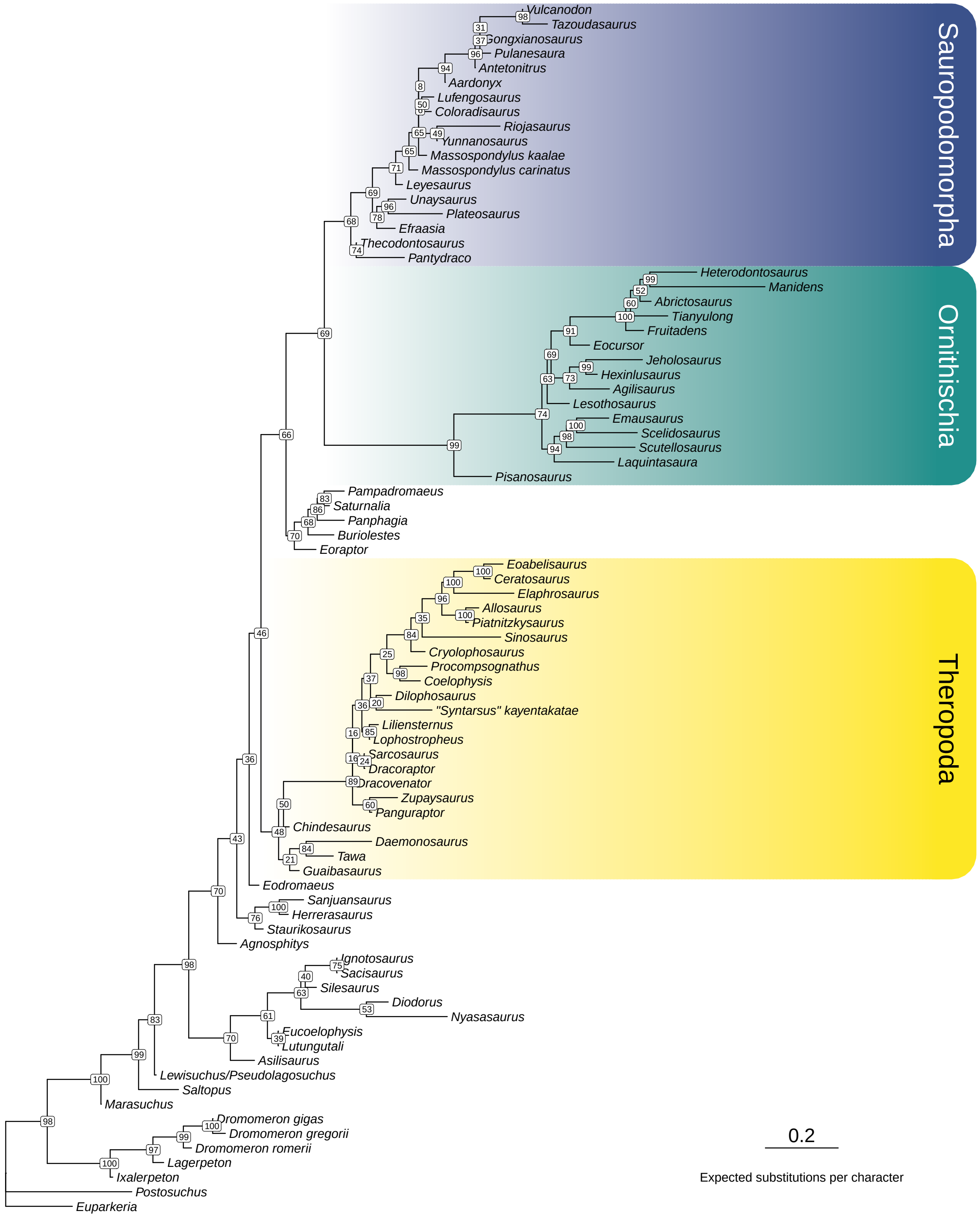


**Supplementary Fig. 14:** Maximum likelihood tree inferred for the LEA matrix without all characters with outlier PS values (characters 206, 318, 169, 391, 198, 338, 377, 306; PS = 2.716–4.454) (ln $L = -6825.859$). Node labels denote ultrafast bootstrap values computed from 1000 replicates.

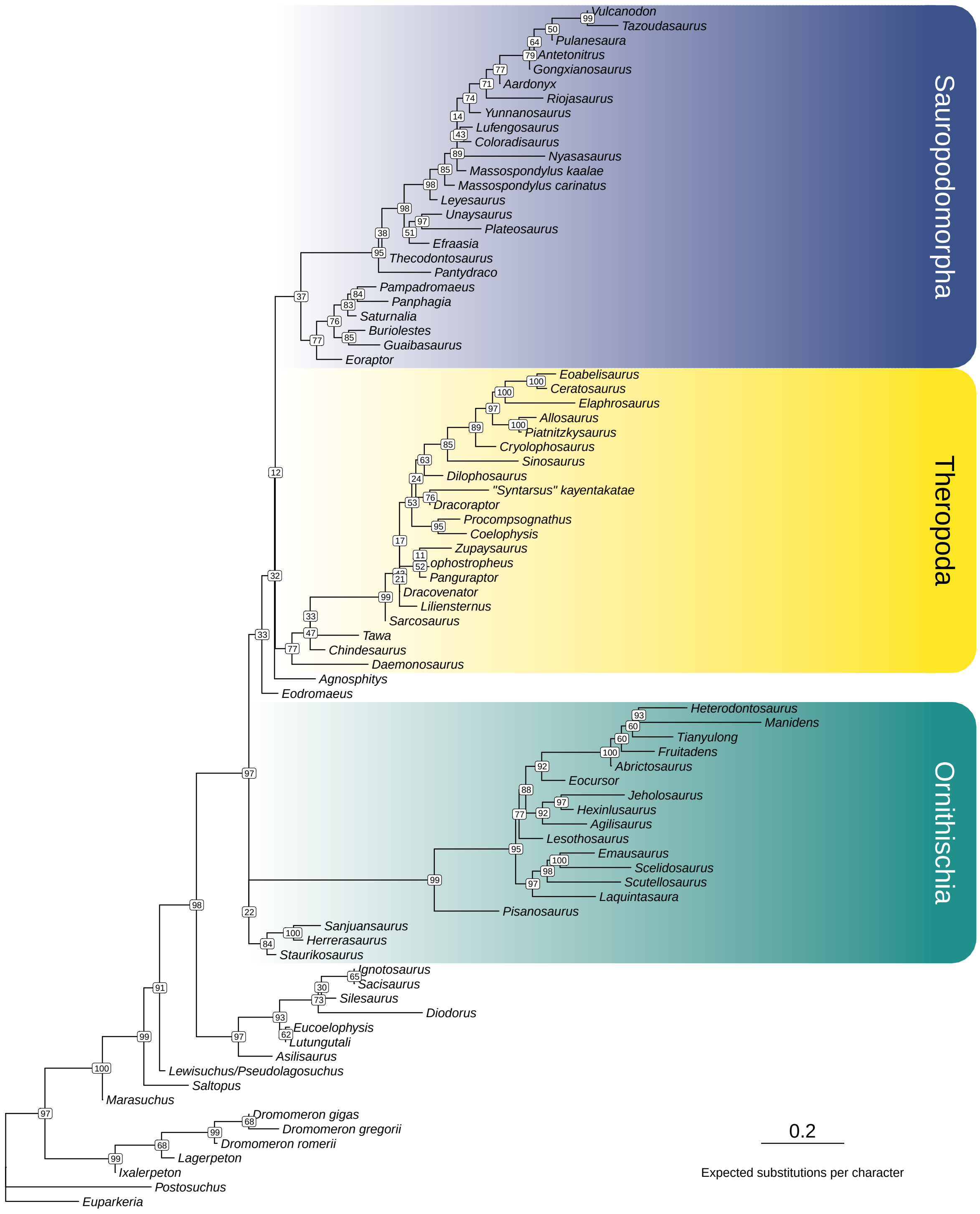


**Supplementary Fig. 15:** Maximum likelihood tree inferred for the LEA matrix with character 77 reverted to its original coding in the BEA matrix ($\ln L = -7013.107$). Node labels denote ultrafast bootstrap values computed from 1000 replicates.

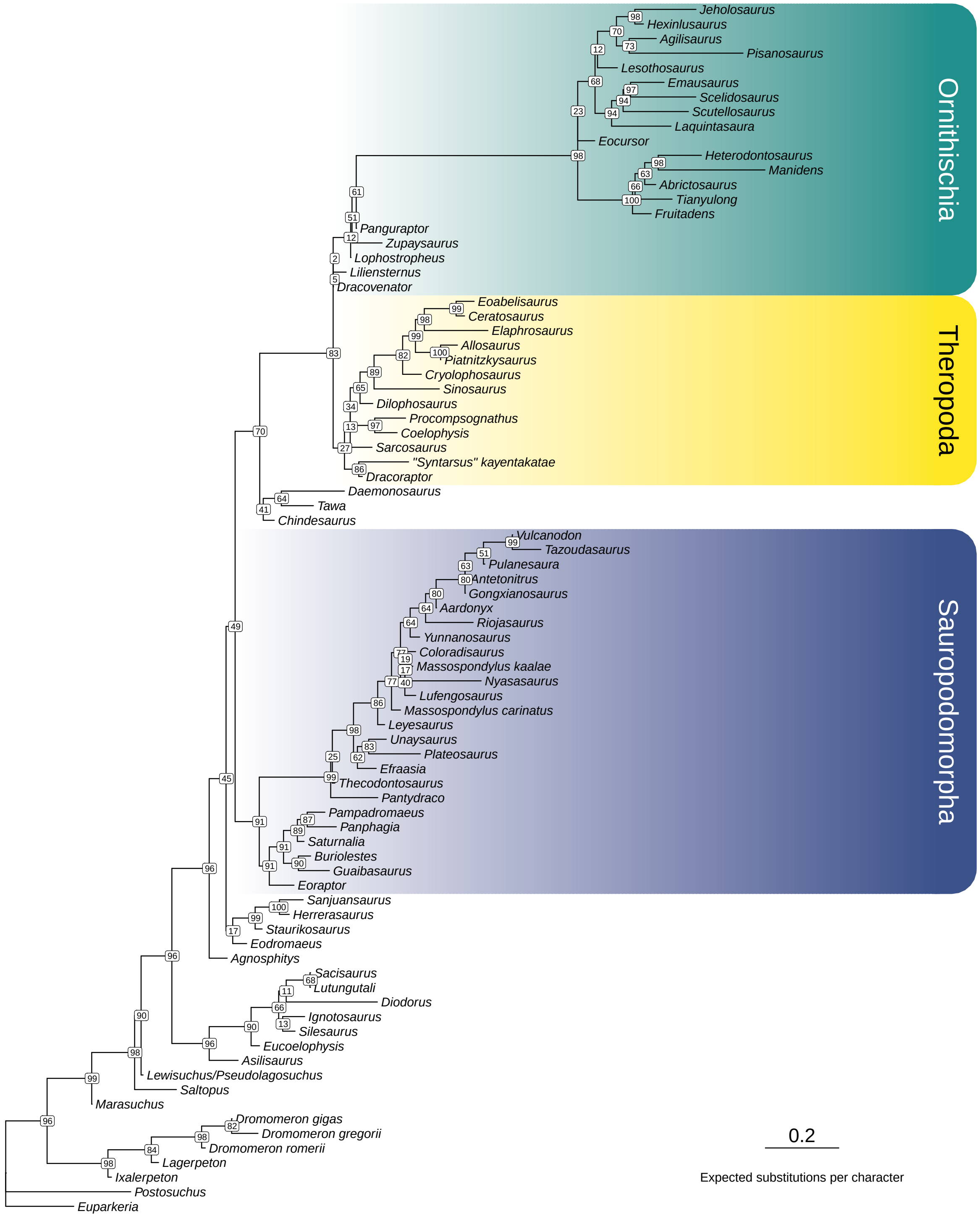


**Supplementary Fig. 16:** Maximum likelihood tree inferred for the LEA matrix with character 148 reverted to its original coding in the BEA matrix ($\ln L = -7010.895$). Node labels denote ultrafast bootstrap values computed from 1000 replicates.

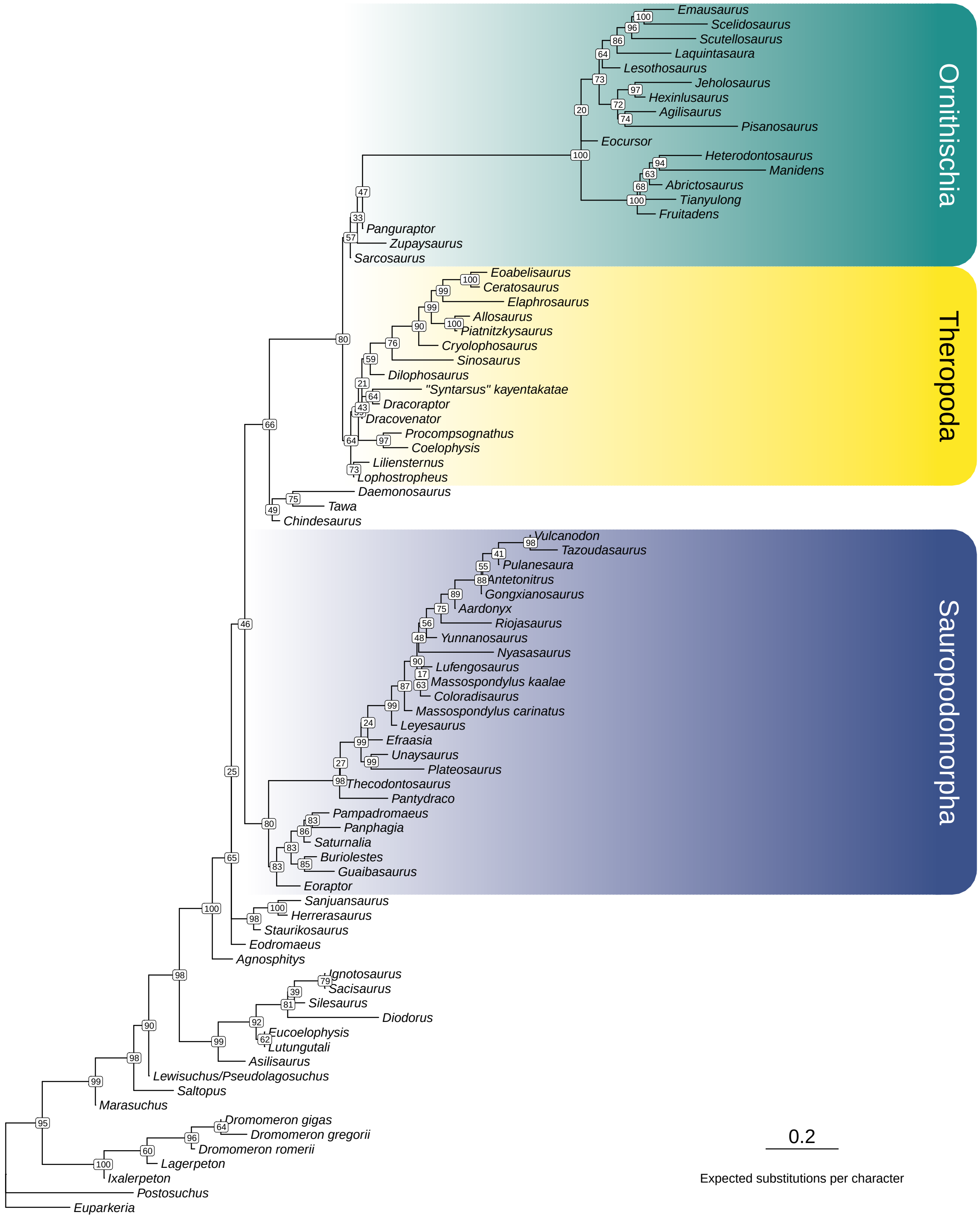


**Supplementary Fig. 17:** Maximum likelihood tree inferred for the LEA matrix with character 363 reverted to its original coding in the BEA matrix (ln $L = -7012.255$). Node labels denote ultrafast bootstrap values computed from 1000 replicates.

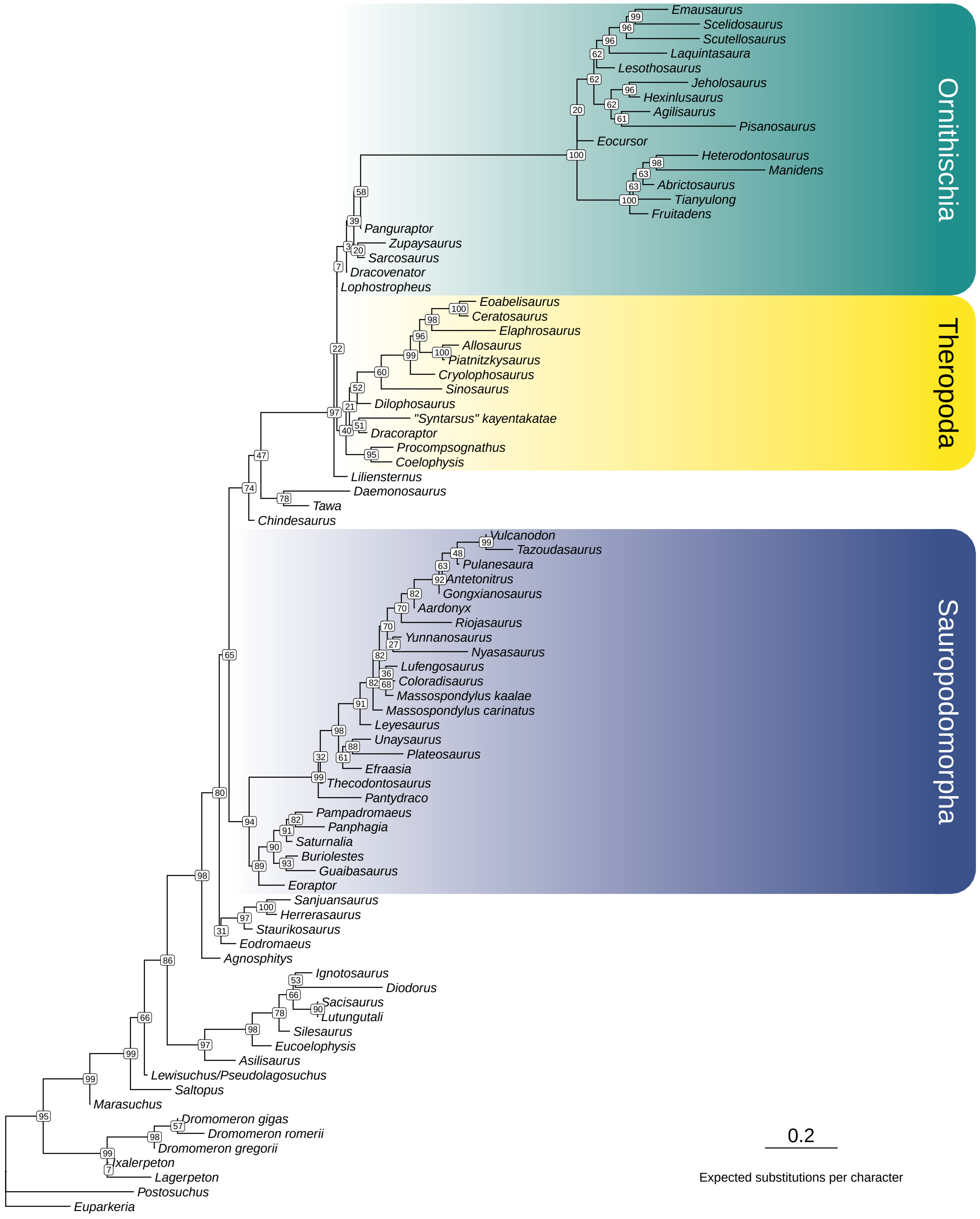


**Supplementary Fig. 18:** Maximum likelihood tree inferred for the LEA matrix with character 370 reverted to its original coding in the BEA matrix ($\ln L = -7023.812$). Node labels denote ultrafast bootstrap values computed from 1000 replicates.

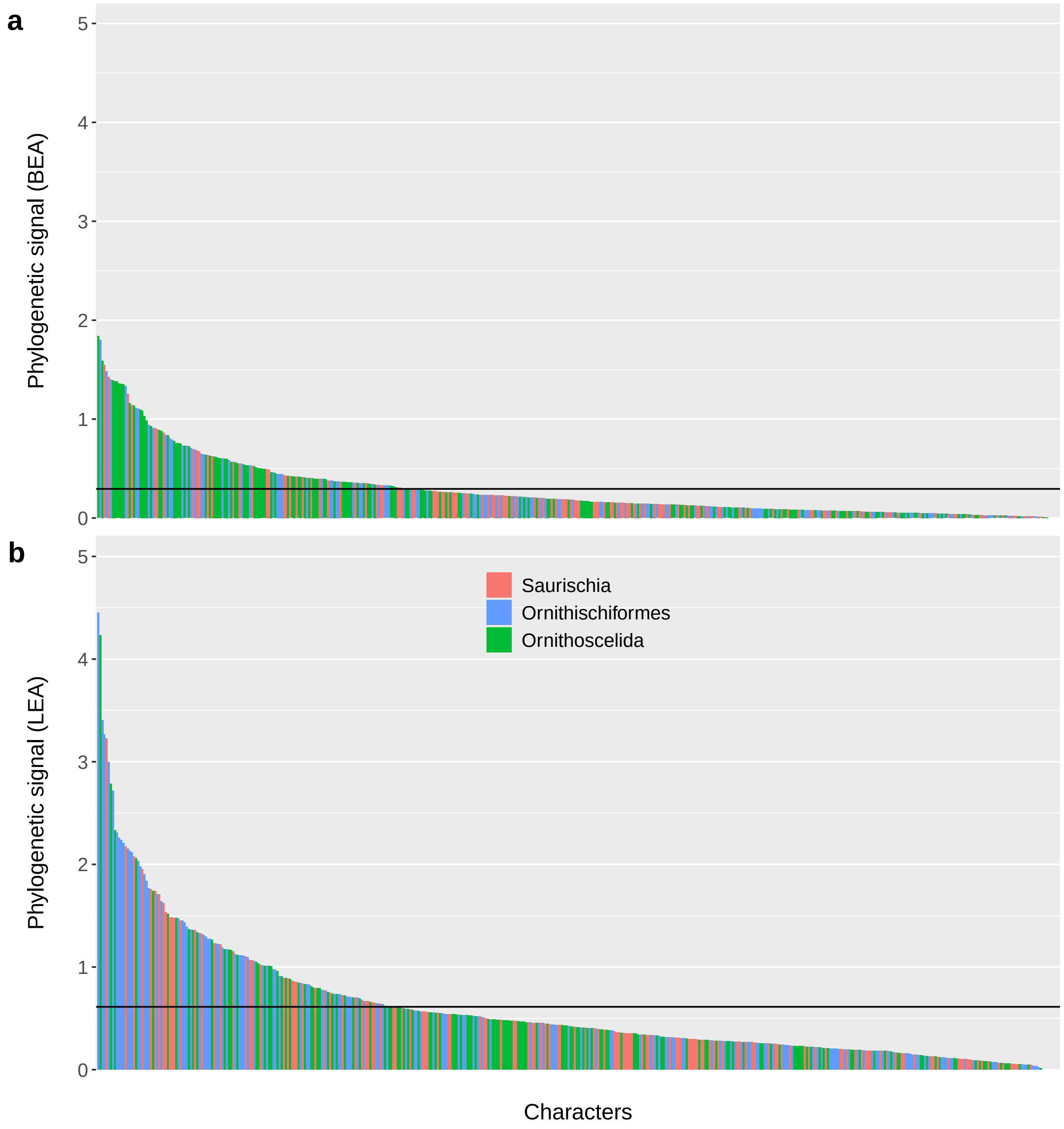


**Supplementary Fig. 19:** Distribution of phylogenetic signal (PS) across the (a) BEA and (b) LEA matrices. The figure corresponds to panels (a) and (c) of Fig. 2 in the main text, except that individual characters are arranged in descending order of their PS values and a common scale is imposed for both datasets. Additionally, the mean PS value in each dataset is denoted by a horizontal line.

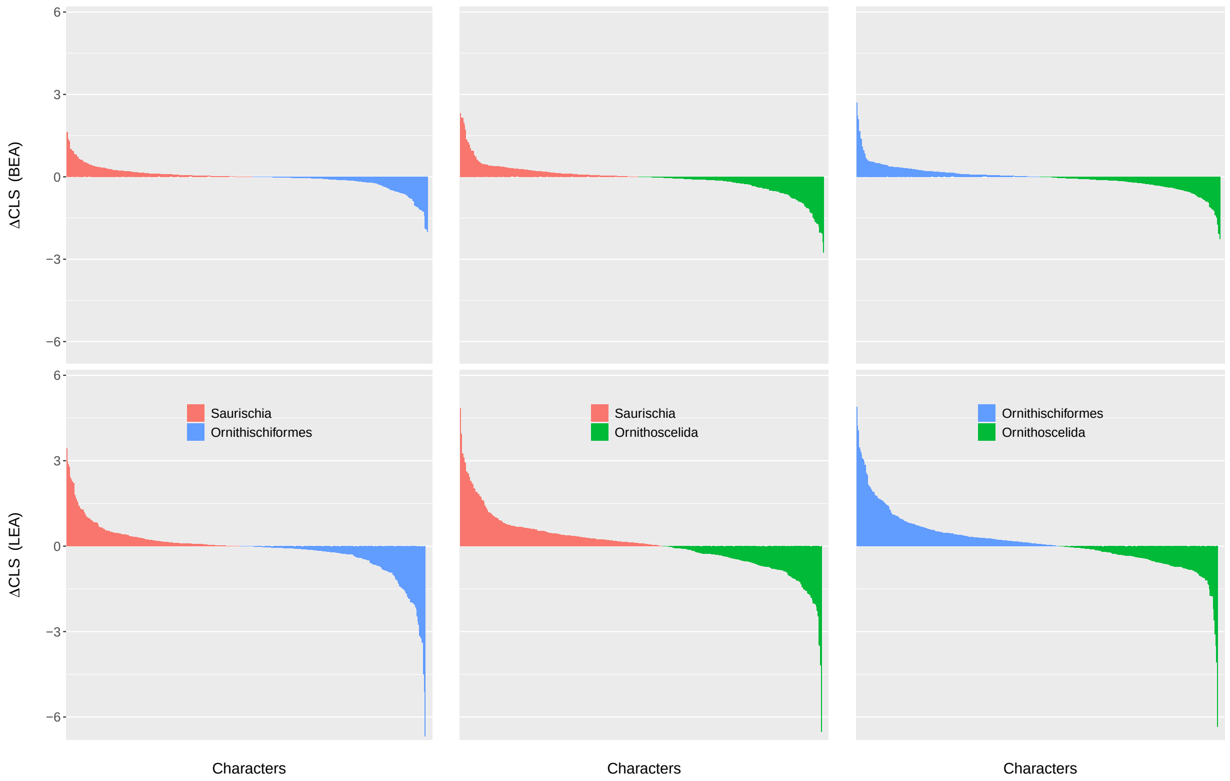


**Supplementary Fig. 20:** Pairwise comparisons of character support for the three alternative early dinosaur topologies. The figure corresponds to Fig. 3 in the main text, except that individual characters are arranged in descending order of their ΔCLS values and a common scale is imposed for both datasets.
